# Multiphasic Organization and Differential Dynamics of Proteins Within Protein-DNA Biomolecular Condensates

**DOI:** 10.1101/2025.06.09.658691

**Authors:** Ashish Shyam Tangade, Anupam Mondal, Jiahui Wang, Young C. Kim, Jeetain Mittal

## Abstract

Biomolecular condensates formed through liquid-liquid phase separation are increasingly recognized as critical regulators of genome organization and gene expression. While the role of proteins in driving phase separation is well-established, how DNA modulates the structure and dynamics of protein-DNA condensates remains less well understood. Here, we employ a minimalist coarse-grained model to investigate the interplay between homotypic protein-protein and heterotypic protein-DNA interactions in governing condensate formation, composition, and internal dynamics. Our simulations reveal that DNA chain length and flexibility critically influence condensate morphology, leading to the emergence of multiphasic and core-shell organizations under strong heterotypic interactions. We find that DNA recruitment into the condensate significantly alters protein mobility, giving rise to differential dynamics of proteins within the condensate. By analyzing the distribution profiles of protein displacements, we identify up to five distinct diffusion modes, including proteins bound to DNA, confined within the dense phase, or freely diffusing. These results provide a mechanistic framework for interpreting spatially heterogeneous protein dynamics observed in chromatin condensates and emphasize the direct role of DNA in tuning condensate properties. Our findings provide new insights into how biophysical parameters may control the functional architecture of protein-DNA condensates in biological systems.

**TOC Graphic:** 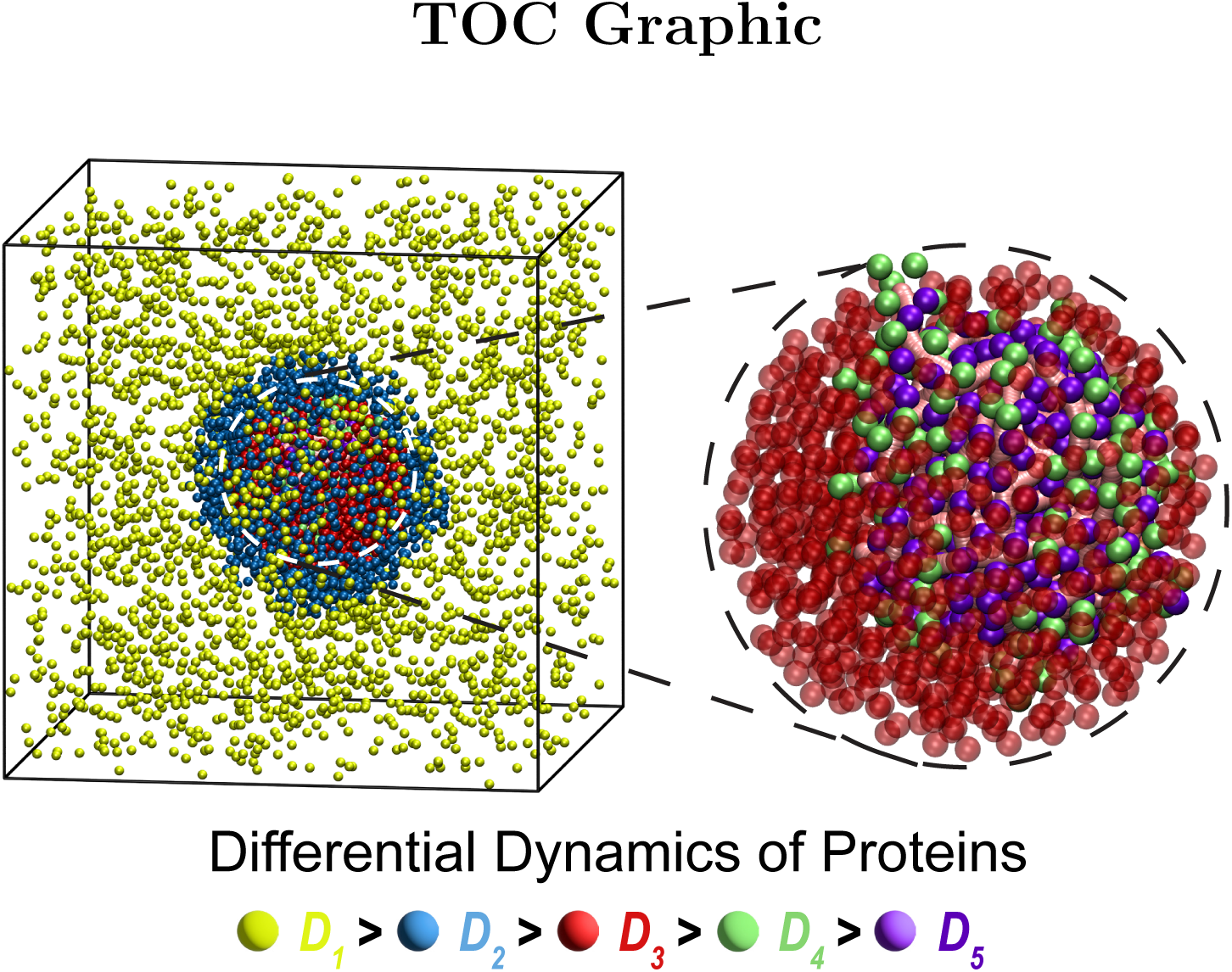

## Introduction

Biomolecular condensates (BMCs), also known as membraneless organelles, are dynamic assemblies in cells formed via liquid-liquid phase separation (LLPS) of proteins and nucleic acids. These condensates lack a surrounding membrane yet maintain a localized high concentration of specific biomolecules that facilitates diverse biological functions.^1,2^ A few common examples of BMCs include nucleolus, stress granules, paraspeckles, Cajal bodies, and mitochondrial nucleoids.^3,4^ BMCs are known to play a crucial role in genome organization, transcriptional regulation, RNA processing, cell signaling, and many more.^5–9^ The assembly and internal organization of BMCs are primarily governed by homotypic interactions between proteins, heterotypic interactions between proteins and nucleic acids, as well as among nucleic acids themselves. These interactions arise from a combination of electrostatics, hydrogen bonding, *π* – *π* stacking, cation–*π* interactions, and hydrophobic effects^10–14^ whose balance determines not only whether phase separation occurs, but also the internal structure, morphology, dynamics, and functional compartmentalization within the condensates.

Several experimental and computational studies have probed the role of protein-DNA interactions in driving phase separation. ^15–24^ For instance, the protein Fused-in-Sarcoma (FUS), which comprises both prion-like and RNA/DNA-binding domains, undergoes phase separation with single- and double-stranded DNA,^17^ revealing that electrostatic and *π*-stacking interactions between FUS and nucleic acids can drive heterotypic condensation. ^19,25,26^ Similarly, the transcription factor FoxA1 forms condensates with DNA and can spatially organize distal DNA regions into close proximity to carry out cellular functions. ^24^ Heterochromatin protein 1 (HP1*α*) is another example that has been shown to compact DNA and form condensates possibly due to homotypic HP1*α*-HP1*α* oligemerization.^27^

In addition to these molecular interactions, a few other experimental studies^27–29^ have recently shown that the physical characteristics of DNA such as its length and flexibility, play critical roles in modulating phase behavior of biomolecular condensates. For example, Ryu *et al.*^28^ demonstrated that the formation of protein-DNA cluster by SMC proteins exhibits a critical dependence on DNA length: for shorter DNA chains, the cluster size remains largely insensitive to DNA lengths, while for longer DNA chains, the cluster size increases strongly with DNA length, thereby promoting phase separation. Likewise, Shakya and King^29^ showed that flexible DNA leads to denser and more compact coacervates with poly-L-lysine, indicating that DNA stiffness influences condensate architecture and internal mobility.

Computational studies using coarse-grained (CG) models^30,31^ have been powerful in exploring protein-DNA condensates across large system sizes and time scales that are inaccessible to all-atom molecular dynamics. Using such CG models, recently it has been shown that the HP1*α* and histone H1 proteins can form multiphasic condensates with layered organization that is mediated by DNA.^32^ Ancona and Brackley^33^ used simple CG simulations to map the phase behavior of HP1*α*–chromatin systems under different interaction regimes, revealing that the relative strength of homotypic versus heterotypic interactions governs phase stability and internal organization. Multiscale simulations have further shown how post-translational modifications, such as phosphorylation of HP1*α*, alter its interaction with DNA and modulate condensate formation and physical properties.^22,23^ Despite these advances, a detailed mechanistic understanding of how homotypic and heterotypic interactions interplay with DNA’s physical properties to determine condensate architecture and protein dynamics remains incomplete.

To address this gap, we develop a minimalistic coarse-grained model to disentangle the roles of homotypic and heterotypic interactions, and DNA physical properties in the formation and organization of protein-DNA condensates. In our model, protein molecules are treated as single beads, while DNA molecule is modeled as a polymer chain with a monomer representing one base pair. By varying protein-protein (homotypic) and protein-DNA (heterotypic) interaction strengths, and different physical properties of DNA chains, we systematically map the phase behavior and structural organization of the resulting condensates.

Our results reveal a rich landscape of condensate behavior. We identify regimes where homotypic interactions alone are insufficient to induce phase separation, but heterotypic interactions with DNA enable condensate formation. Strikingly, when both interaction types are present, condensates adopt a multiphasic architecture, characterized by DNA-bound protein-rich cores surrounded by a protein-enriched shell. Furthermore, we show that DNA length and flexibility critically modulate the phase separation and the internal organization of condensates. Importantly, we uncover distinct dynamical modes of proteins within condensates, including freely diffusing, transiently DNA-bound, and stably bound populations, suggesting spatially heterogeneous molecular mobility. Collectively, our results reveal how the interplay between interaction types and DNA physical features shapes the structure, composition, and dynamic behavior of protein-DNA condensates, providing mechanistic insight into principles underlying nuclear organization.

## Materials and Methods

Simulating large-scale phenomena such as LLPS requires careful balance of computational expense with system size. In this work, we develop a minimalistic coarse-grained (CG) model of protein and DNA and their interactions to study the phase separation of biomolecular condensates. Similar CG models have been widely used to study intrinsically disordered proteins and their phase behavior, ^34–39^ as well as protein–DNA complexes and chromatin organization with great success.^22,32,33,40^

### Protein Model

Protein molecule is modeled as a single spherical bead, as schematically shown in Fig. 1A. The size of the protein bead is considered based on the radius of gyration (*R_g_*) of DNA-binding proteins typically implicated in biomolecular condensates. Specifically, we set the diameter of the protein bead to *σ_P_* = 50 Å, which approximates twice the average *R_g_* value for proteins such as TFAM and HP1*α*.^22,41^ The mass of each protein bead is chosen as 5000 g/mol.

**Figure 1:**
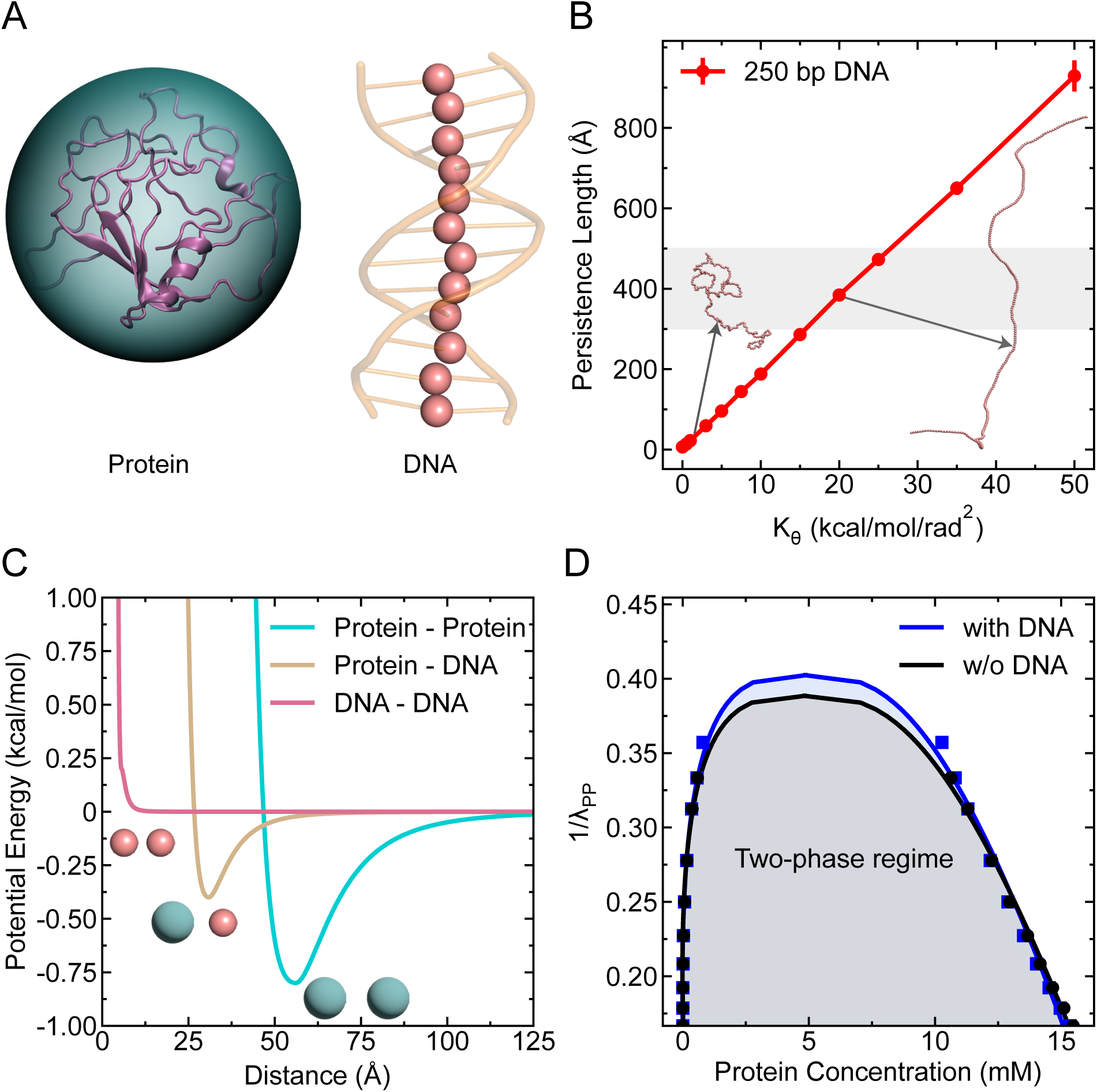
Description of the minimalistic model. (A) Schematic of the CG model: protein is modeled as a single spherical bead, and DNA as bead-spring polymers with each bead representing one base pair. (B) Persistence length of the polymeric DNA of length 250 bp as a function of the angle potential parameter *K_θ_*. The shaded region indicates the experimentally observed range of persistence length for dsDNA. Representative DNA conformations are shown for two *K_θ_* values. (C) Ashbaugh-Hatch potential energy for protein-protein (homotypic), protein-DNA (heterotypic), and DNA-DNA interactions as a function of inter-bead distance. (D) Phase diagrams showing protein phase separation as a function of homotypic interaction strength *λ_PP_*, in the absence and presence of DNA (*λ_PD_* = 0.5, DNA length *L* = 250 bp), exhibiting phase coexistence.

### DNA Model

The DNA molecule is modeled as a polymeric chain in which each monomer corresponds to one base pair (bp), as schematically illustrated in Fig. 1A. The total potential energy of the DNA chain comprises bond stretching and angular bending contributions, as given by

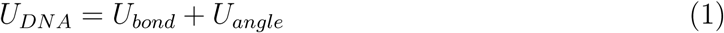

The bond energy between successive base pairs is given by a harmonic potential:

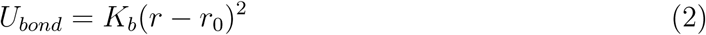

where *K_b_*= 20 kcal/mol/Å^2^ is the spring constant and *r*_0_ = 5.5 Å is the equilibrium bond length.

To modulate the bending rigidity of DNA, the angular potential between three consecutive monomers is defined as:

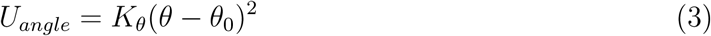

where *K_θ_* is the spring constant for angle potential and *θ*_0_ = 180*^◦^* is the equilibrium angle between two successive bonds of DNA. In order to determine the flexibility of the DNA chain ideal for double-stranded DNA (dsDNA), we varied *K_θ_* values and measured the persistence length (see Supporting Information for calculation details) of a single DNA chain of length 250 bp as shown in Fig. 1B. The experimentally^42,43^ reported values of persistence length for dsDNA is highlighted with grey shaded region, based on which we set the *K_θ_* value to 20 kcal/mol/rad^2^ that qualitatively replicate the flexibility of dsDNA. Additionally, the diameter (*σ_D_*) and mass of each monomer are chosen as 5 Å and 500 g/mol respectively.

### Protein-DNA Interactions

For modeling of the nonbonded interactions between protein and DNA molecules, we used a modified Lennard-Jones (LJ) potential,^34,44^ where the attractive interactions between two particles are scaled independently of the short-range repulsive interactions. The nonbonded interactions are modeled by the following Ashbaugh-Hatch ^44^ potential

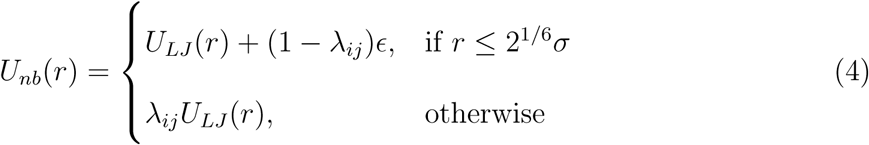

in which *U_LJ_*(*r*) is the standard Lennard-Jones potential,

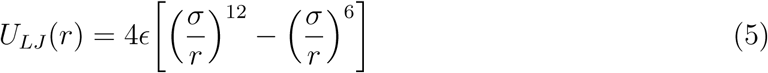

where *ɛ* is the interaction strength fixed to a value of 0.2 kcal/mol and *λ_ij_* is the hydropathy between two interacting species. The arithmetic average is set as the combination rule for the size *σ* (i.e., *σ_PD_* = (*σ_P_* + *σ_D_*)*/*2). The strength of homotypic and heterotypic interactions can be varied by changing the hydropathy value *λ* (Fig. 1C), while the interaction between DNA molecules is made repulsive by setting *λ_DD_* = −1.

### Simulation Protocol

To explore the phase behavior of protein–DNA mixtures, we simulated systems with a fixed stoichiometric ratio of protein to DNA monomers as 1:1. Specifically, we considered 5000 CG protein beads and a number of DNA chains of a given length *L* base pairs such that the total number of DNA beads also equaled 5000. All particles were initially placed randomly at the center of a cubic box with an edge length of 2000 Å with periodic boundary conditions in all directions. We performed Langevin dynamics simulations at a fixed temperature of 300 K, with the friction coefficient *γ_i_* = *m_i_/t*_damp_, where *m_i_* is the mass of the *i*^th^ particle and *t*_damp_ is the damping factor that is set to 1000 ps. The equations of motion were integrated using a velocity-Verlet algorithm with a time step of 10 fs. All simulations were performed using the HOOMD-blue molecular dynamics engine (version 2.9.3),^45,46^ utilizing the additional features provided by azplugins (version 0.11.0).

## Results and Discussions

To illustrate the effectiveness of our minimal model in capturing biomolecular condensate formation, we first examined whether it could reproduce phase separation in systems with only proteins and in mixed systems containing both proteins and DNA. For this, we varied the interaction strength *λ_PP_* between proteins in the absence and presence of DNA and performed 500 ns simulations for each *λ_PP_* value. We divided the simulation trajectory into five equal blocks in which the first block (i.e., initial 20% of the data) was discarded for equilibration and computed the mean and standard error by calculating the block average from the remaining data. As shown in Fig. 1D, both systems exhibit clear signatures of coexistence of two phases, characterized by the emergence of a dense condensate coexisting with a surrounding dilute phase, in which most DNA chains are recruited into the condensate and lower fraction of proteins stay in the dilute phase for the protein-DNA system. This result suggests that our model is well suited to study condensate formation and gives us the confidence to use further in dissecting how homotypic (protein–protein) and heterotypic (protein–DNA) interactions influence the phase behavior of protein–DNA condensates.

### Phase Behavior of Protein-DNA Condensates as a Function of Homotypic and Heterotypic Interactions

To delineate the role of homotypic and heterotypic interactions in the formation and composition of protein-DNA condensates, we systematically varied the homotypic (*λ_PP_*) and heterotypic (*λ_PD_*) interaction strengths across a broad range of parameter space (Fig. 2).

**Figure 2:**
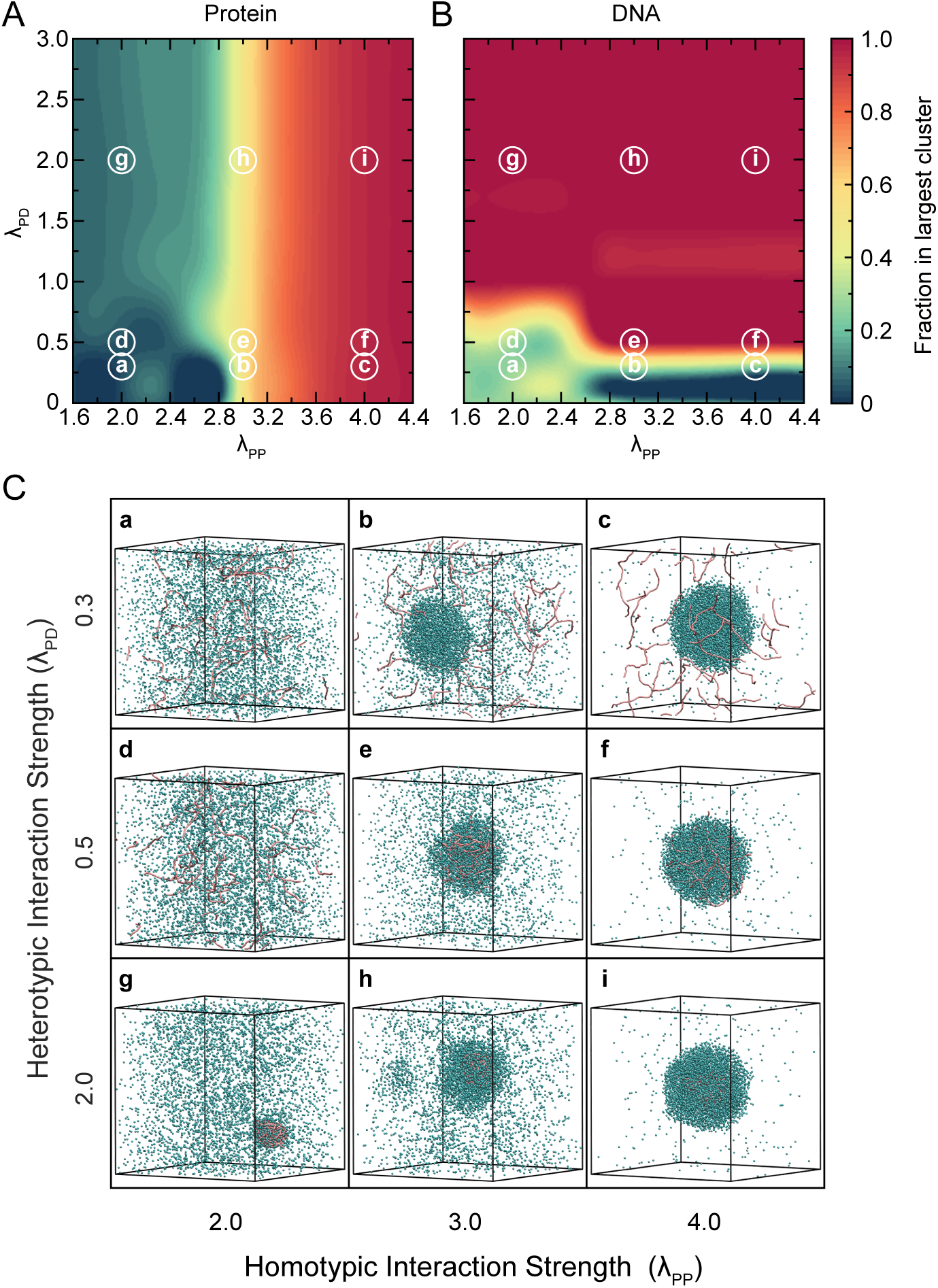
Phase space of homotypic and heterotypic interactions. Heatmap showing the average fraction of (A) protein and (B) DNA in the largest cluster for varying homotypic (*λ_PP_*) and heterotypic (*λ_PD_*) interaction strengths. (C) Representative snapshots from simulation for different regimes of homotypic and heterotypic interactions, labeled (a–i). Small letters circled in (A) and (B) correspond to the respective subfigures shown in (C). Proteins are shown as cyan beads whereas DNA molecules are represented by pink colored chains. Each snapshot illustrates distinct condensate compositions: (a, d) no condensation, (b, c) protein-only condensates, (e, f) mixed protein-DNA condensates with fluid-like DNA, (g) DNA-scaffolded protein clusters via bridging, and (h, i) dense protein-DNA co-condensates with highly localized DNA cores. DNA chains of length 250 bp (20 in total) and 5000 protein beads are used in these simulations.

For these simulations, we used DNA chains of length 250 bp with 20 such chains present in the system. We performed 2 *µ*s long simulation for any combination of *λ_PP_* and *λ_PD_* in which the initial 400 ns of each trajectory was discarded as equilibration, and the remaining 1.6 *µ*s of data was used for analysis. To quantify condensate formation, we evaluated the protein fraction (Fig. 2A) and DNA fraction (Fig. 2B) within the largest cluster. These fractions provide insight into both the size and composition of the condensates (whether they are protein-only, DNA-only, or mixed protein–DNA) and directly reflect how the interaction strengths drive phase separation. For the calculation of the largest cluster, we used a distance criterion where molecules within 1.5 *σ_PD_* are considered as part of the same cluster using the Freud^47^ Python library (version 2.12.1).

The resulting phase diagrams (Fig. 2A,B) reveal a diverse landscape of condensate behaviors, characterized by distinct regimes determined by the interplay between homotypic and heterotypic interactions. For instance, when the homotypic interactions are strong (*λ_PP_* ≥ 3) and the heterotypic interaction is weak (*λ_PD_ <* 0.5), we observe protein-only condensates in which proteins undergo phase separation independent of DNA. In this regime, DNA molecules remain dispersed in the dilute phase and are excluded from the condensate (Fig. 2C, insets b and c). This organization of proteins undergoing phase separation independent of DNA is observed in *in vitro* experiments of FUS protein, where the low complexity domain of FUS is responsible for interactions causing phase separation.^17,48,49^ With protein Ddx4, similar arrangement of components is observed where protein forms a condensate via homotypic interactions but does not recruit dsDNA in the condensate. ^50^ In contrast, when both the homotypic and heterotypic interaction strengths are weak (*λ_PP_ <* 3 and *λ_PD_ <* 0.5), neither proteins nor DNA undergo phase separation, and the system remains homogeneous with all components remaining dispersed in the dilute phase (Fig. 2C, inset a). This regime represents a baseline state where interaction strengths are insufficient to drive condensate formation. As the heterotypic interaction strength increases to a moderate range (0.5 ≤ *λ_PD_* ≤ 1.0) while maintaining strong homotypic interactions, both proteins and DNA are recruited into the condensate, forming a mixed protein-DNA co-condensate. In this regime, the DNA remains fluid-like and dispersed across the entire condensate (Fig. 2C, insets e and f). This organization of fluidized DNA can be of functional relevance for the transcriptional condensates, where the DNA needs to be accessible to transcriptional machinery, enhancers and co-activators.^51^ Interestingly, when heterotypic interactions are strong (*λ_PD_ >* 1.0) but homotypic interactions are weak (*λ_PP_ <* 3), DNA molecules serve as scaffolds and proteins alone do not phase separate but instead become localized by binding to DNA, forming “bridging” contacts between different DNA segments. This results in localized protein clusters around DNA with a relatively low protein fraction in the overall condensate (Fig. 2C, inset g). This condensate formation is similar to the bridging induced phase separation which has been proposed as another mechanism for phase separation of protein-DNA condensates ^21,40,52^ and has also been experimentally reported in condensates formed with cohesion-SMC protein complexes^28^ and DNA-binding proteins from starved cells.^53^ Furthermore, when both homotypic and heterotypic interactions are strong, proteins undergo phase separation and simultaneously form bridging contacts with DNA. This leads to compact, dense co-condensates where DNA is sequestered in a highly localized region near the center of the droplet (Fig. 2C, insets h and i). Such an arrangement within condensate is like the ones reported for the HP1*α*-DNA system where proteins are dynamic, but the DNA molecules are localized within a smaller region in the condensate.^27^ It also aligns with the hierarchical condensation observed in the protein Krüppel-like factor 4 (Klf4) on DNA, where Klf4 forms an adsorbed layer on DNA at lower protein concentrations and these adsorbed layers further form thicker condensate at higher protein concentrations.^54^

Based on these observations, we categorize the heterotypic interaction strength into three regimes: weak (*λ_PD_ <* 0.5), moderate (0.5 ≤ *λ_PD_* ≤ 1.0), and strong (*λ_PD_ >* 1.0), and the homotypic interaction into two regimes: weak (*λ_PP_ <* 3) and strong (*λ_PP_* ≥ 3). This classification helps us to further evaluate the impact of DNA physical features for modulating condensate properties within each interaction regime.

### Role of Heterotypic Interactions in the Organization of Protein-DNA Condensates

We next explored how homotypic and heterotypic interactions govern the internal organization of the components within the condensate. Specifically, we focused on two key aspects: the spatial configuration of DNA chains and the extent of protein binding to DNA, both of which are critical for understanding the structural and functional implications of condensate formation.

To probe DNA organization within the condensate, we computed the average radius of gyration (*R_g_*) of DNA molecules (see Supporting Information for calculation details) as a function of protein-DNA interaction strength (*λ_PD_*), for three different homotypic proteinprotein interaction strengths (*λ_PP_* = 2.0, 3.0, 4.0). As shown in Fig. 3A, when protein-DNA interaction strength is weak (*λ_PD_ <* 0.5), the DNA chains remain extended, exhibiting an average *R_g_* of approximately 295 Å which is comparable to the value obtained for an isolated DNA chain of the same length, as indicated by the black dashed line. This suggests that under weak protein-DNA interactions, DNA adopts a relaxed conformation. As *λ_PD_* increases, DNA becomes increasingly compact, with *R_g_* dropping to as low as 100 − 125 Å. This compaction is most pronounced when the protein-protein interaction is weak (*λ_PP_* = 2.0), meaning proteins cannot phase separate on their own and instead use DNA as a scaffold to assemble. This compaction behavior closely resembles that of protamine proteins, which causes extreme DNA compaction in sperm cells potentially rendering DNA transcriptionally inactive.^55^ Interestingly, dephosphorylation of protamine,^56^ which have stronger interactions with DNA, show even higher DNA compaction which is in accordance with the trend observed in our result. However, as we increase the homotypic interactions to *λ_PP_* = 3.0 and 4.0, the DNA becomes less compact (larger *R_g_*) for the same *λ_PD_*. This suggests a competition between protein-protein and protein-DNA interactions: when proteins strongly interact with each other, fewer proteins are available to bind and compact DNA. As a result, DNA adopts a more expanded configuration. Such an expansion of DNA in the presence of strong protein-protein interactions might play a significant role in accessibility of DNA to transcription assemblies, cofactors, and activators which then helps DNA to perform its function with these additional assemblies.^18,51,57^

**Figure 3:**
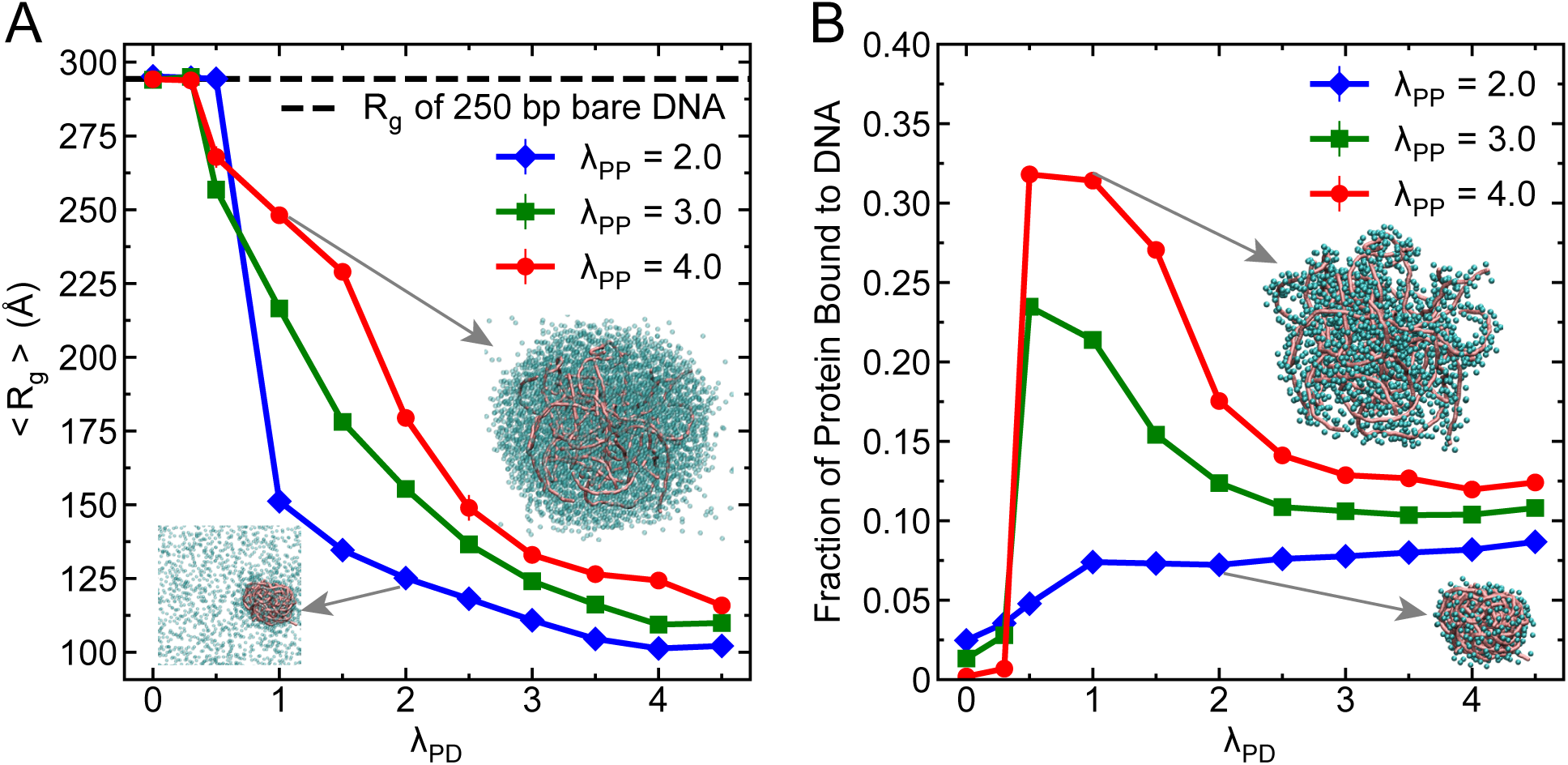
Heterotypic interactions affect the DNA compaction and protein bound to DNA. (A) Average radius of gyration (*R_g_*) of 250 bp DNA chains as a function of heterotypic protein-DNA interaction strength (*λ_PD_*) for three different homotypic proteinprotein interaction strengths *λ_PP_* . The horizontal dashed line indicates the *R_g_* of a single DNA chain of the same length of 250 bp in the absence of proteins. Two representative snapshots (insets) illustrate the structural organization of DNA within the condensate under weak and strong homotypic interactions. (B) Fraction of proteins bound to DNA as a function of *λ_PD_*, for the same three values of *λ_PP_* . Two representative snapshots (insets) show protein binding patterns in the condensates for weak and strong homotypic interactions. DNA molecules are represented in pink color and protein molecules are represented in cyan color. In both panels, error bars represent standard errors obtained from block averaging and the associated error bars are smaller than the symbol size.

For evaluating the organization of proteins within the condensate, we calculated the fraction of proteins bound to DNA using a distance-based criterion where a protein within distance of *σ_P_* + *σ_D_* is considered as bound to DNA. Fig. 3B shows the dependence of this bound fraction on *λ_PD_* for different *λ_PP_* values. With increasing heterotypic interaction strength one would expect that more proteins will bind to DNA, but this is only true for the case when DNAs are acting as a scaffold for proteins (*λ_PP_* = 2.0) (Fig. 3B). Whereas with increasing homotypic interactions (*λ_PP_* = 3.0, *λ_PP_* = 4.0), as proteins form condensate, the trend shows non-monotonic behavior with increasing heterotypic interaction strength: fraction of bound proteins initially increases with *λ_PD_*, peaks at intermediate values (*λ_PD_* ≤ 1.0) and then decreases and saturates at higher heterotypic interaction strengths. This non-monotonic behavior can be explained by considering the effect of DNA compaction. As *λ_PD_* increases, DNA becomes more compact (as seen in Fig. 3A), which reduces the surface area available for proteins to bind (see snapshots in the inset of Fig. 3B). At the same time, proteins are already forming condensates due to strong homotypic interactions, making it harder for them to access tightly packed DNA.

Taken together, both the organization of DNA and the extent of protein-DNA binding within condensates are jointly controlled by homotypic and heterotypic interaction strengths. Even when heterotypic interactions are strong, DNA can become less accessible due to compaction or competition from protein-protein homotypic interactions. These results highlight the complex interplay between protein and DNA interactions in determining the structure and function of biomolecular condensates.

### Role of DNA Length in the Organization of Protein-DNA Condensates

Next, we turn to investigate how the physical properties of DNA affect the structural organization and internal architecture of protein-DNA biomolecular condensates. For this, we first explored the role of DNA chain length in modulating the condensate organization and how the heterotypic and homotypic interactions act upon DNA molecules of varying lengths. Previous experimental studies have shown that longer DNA molecules facilitate protein phase separation by reducing the saturation concentration of proteins for phase separation and form morphologically different structures of condensates.^27^ Additionally, Ryu *et al.*^28^ have shown that condensates formed by bridging-induced phase separation have a critical dependency of DNA length. Despite these insights, the different assemblies of a condensate generated due to DNA length remain less explored. To address this, we systematically varied the DNA length across a wide range of 10, 50, 100, 250, 500, and 1000 base pairs while maintaining a constant heterotypic interaction strength at *λ_PD_* = 2.0 and explored the influence of three different homotypic interaction strengths of *λ_PP_* = 2.0, 3.0, 4.0. This setup allows us to assess how DNA length, in concert with molecular interactions, governs condensate organization.

To quantify DNA organization within the condensate, we computed the average *R_g_* of DNA chains and normalized it by the *R_g_* of the same DNA chains in the absence of any attractive interactions i.e., when both *λ_PP_* and *λ_PD_* are zero. This *R_g_* ratio serves as a measure of DNA compaction induced by attractive protein interactions and the result is shown in Fig. 4A as a function of DNA length. We observe that for shorter length of DNA (10 and 50 bps) whose persistence length is less than the DNA contour length, these molecules act as rigid rods and undergo negligible compaction for all *λ_PP_* values. In contrast, longer DNA molecules (250–1000 bps) show significant compaction, with the degree of compaction depends on the strength of protein-protein interactions. An intriguing trend emerges at an intermediate DNA length of 100 bp, where we observe a pronounced peak in the *R_g_* ratio for *λ_PP_* = 3.0 and 4.0. This suggests that rather than compacting, DNA chains at this length adopt a more expanded and possibly ordered conformation within the condensate (see corresponding snapshot of DNA structures in the inset of Fig. 4A). Such behavior may reflect a structural rearrangement wherein proteins form dense networks around the DNA, leading to steric expansion instead of compaction. Interestingly, across increasing DNA lengths, we find that the strongest DNA compaction generally occurs at the lowest homotypic interaction strength (*λ_PP_* = 2.0), followed by moderate compaction at *λ_PP_* = 3.0, and least compaction at *λ_PP_* = 4.0. This inverse relationship between homotypic interaction strength and DNA compaction at fixed heterotypic strength likely arises because strong protein-protein interactions favor self-association of proteins, reducing their availability to engage and wrap around the DNA, thereby attenuating compaction. To evaluate the robustness of these observations, we performed similar analyses under both moderate (*λ_PD_* = 0.5) and much stronger (*λ_PD_* = 3.5) heterotypic interaction regimes (Figs. S1A and S1C). At high *λ_PD_* = 3.5, the overall trend of stronger compaction at lower *λ_PP_* is preserved, but notably, the peak at 100 bp DNA seen for *λ_PP_* = 3.0, 4.0 vanishes. This indicates that strong protein-DNA interactions lead to uniform compaction across intermediate chain lengths (Fig. S1C). Conversely, under moderate heterotypic interaction strength (*λ_PD_* = 0.5), we observe that DNA compaction is minimal for *λ_PP_* = 2.0 (ratio remains nearly invariant across lengths), while for *λ_PP_* = 3.0 and 4.0, compaction increases modestly beyond 100 bp (Fig. S1A). For the case of *λ_PP_* = 3.0, the result shows relatively higher compaction of DNA than *λ_PP_* = 4.0, possibly reflecting an optimal balance between protein-protein association and DNA engagement in this intermediate interaction regime.

**Figure 4:**
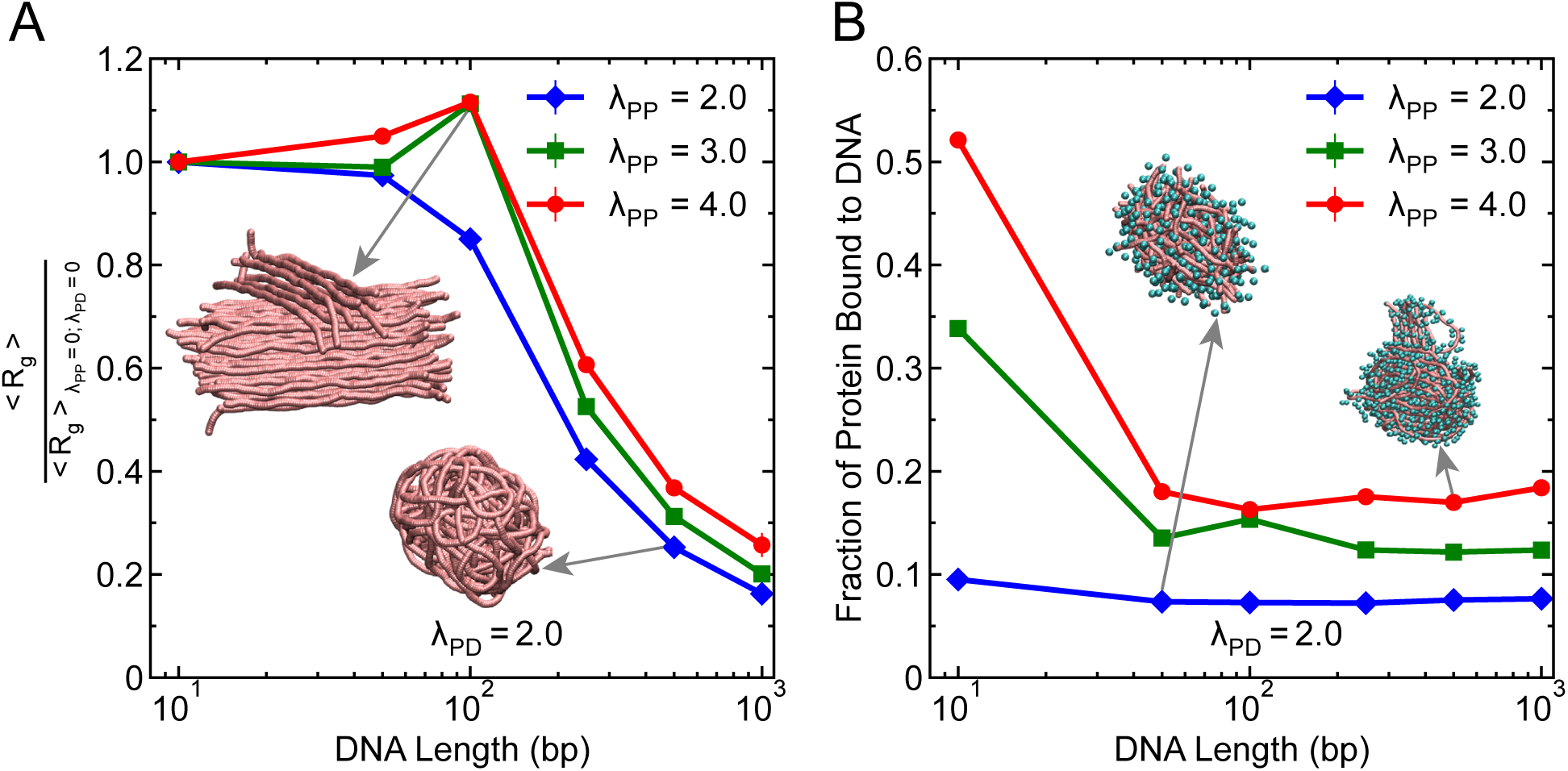
DNA length affects the organization within the condensates. (A) Normalized *R_g_* of DNA within the condensate, plotted as a function of DNA length for three different homotypic interaction strengths (*λ_PP_* = 2.0, 3.0, 4.0) at a fixed heterotypic interaction strength *λ_PD_* = 2.0. *R_g_* is normalized by the *R_g_* of corresponding DNA chains in the absence of attractive interactions (*λ_PP_* = *λ_PD_* = 0). Insets show representative conformations of DNA chains (proteins are removed for clarity) at selected lengths and homotypic strengths. (B) Fraction of proteins bound to DNA within the condensate as a function of DNA length, for the same interaction strengths as in panel (A). Insets show the structural organization of proteins bound with DNA at selected lengths and homotypic strengths. DNA molecules are represented in pink color and protein molecules are represented in cyan color. In both panels, error bars represent standard errors obtained from block averaging and the associated error bars are smaller than the symbol size.

To further elucidate the effect of DNA length on protein organization within the condensate, we computed the fraction of proteins bound to DNA as a function of DNA length (Fig. 4B). Surprisingly, the fraction of bound proteins remains nearly constant across DNA lengths, except for the shortest chain (10 bp), which exhibits the highest protein binding. This elevated binding at 10 bp is likely due to the extended, rod-like conformation of these short chains and this DNA length (approximate contour length is 50 Å) is similar to the diameter of the protein, making their surfaces maximally accessible to proteins. For longer chains, despite their increased contour length and compaction, the overall protein-DNA contact fraction remains invariant. This suggests that once a minimal DNA scaffold is available for bridging, further increases in chain length do not significantly alter the bound protein distribution in the condensate. However, increasing homotypic interaction strength (*λ_PP_*) consistently leads to higher DNA-bound protein fractions for all DNA lengths (Fig. 4B). This trend can be explained by the partial expansion of DNA within the condensate at higher *λ_PP_* (as seen in Fig. 4A), which increases surface exposure and accessibility of DNA (see representative snapshots in the inset), thereby facilitating additional protein binding. The same qualitative trends are reproduced in simulations at moderate and strong heterotypic strengths (Figs. S1B, D).

### Role of DNA Flexibility in the Organization of Protein-DNA Condensates

Next, we further explored the influence of another critical physical parameter, the flexibility of DNA, on the organization of protein-DNA condensates. DNA flexibility is known to differ significantly between single-stranded DNA (ssDNA) and double-stranded DNA (dsDNA), with ssDNA exhibiting a lower persistence length and higher conformational flexibility. Motivated by this difference, we systematically varied DNA flexibility of 250 bp long DNA chains by tuning the angular spring constant *K_θ_* in the bending potential (Eq. 3), which effectively modulates the DNA’s persistence length (Fig. 1B). Specifically, we considered four different *K_θ_* values: 0.5, 1.5, 7.5, and 20 kcal/mol/rad^2^, which are chosen in such a way that 0.5 kcal/mol/rad^2^ models the ssDNA flexibility, 1.5 kcal/mol/rad^2^ represents a semi-flexible DNA, 7.5 kcal/mol/rad^2^ resembles a semi-rigid DNA and 20 kcal/mol/rad^2^ captures the dsDNA flexibility (Fig 1C). Recent experiments suggest that flexible DNA increases the propensity for formation of protein-DNA condensates as the penalty of bending a flexible chain is lower.^29,58,59^ To evaluate how DNA flexibility modulates condensate organization, we computed the average *R_g_* of DNA and the fraction of proteins bound to DNA within the dense phase for different *K_θ_* values at a fixed heterotypic interaction strength *λ_PD_* = 2.0 for three different homotypic interaction strengths of *λ_PP_* = 2.0, 3.0, 4.0.

As shown in Fig. 5A, the average DNA *R_g_* increases monotonically with increasing *K_θ_*, suggesting that more rigid DNA chains maintain a more expanded conformation regardless of homotypic interaction strength. This trend is consistent with expectations from bending rigidity: more rigid chains resist compaction due to the higher energetic cost of deformation. Interestingly, the degree of compaction exhibits a nontrivial dependence on homotypic interaction strength. The most compact conformations are observed at the lowest *λ_PP_* = 2.0, followed by moderate compaction at *λ_PP_* = 3.0, and the least compaction at *λ_PP_* = 4.0, mimicking the similar trend as observed in the case of DNA chain length. To obtain the generality of these results, we performed similar analysis under both moderate (*λ_PD_* = 0.5) and much stronger (*λ_PD_* = 3.5) heterotypic interaction regimes (Figs. S2A and S2C). At strong *λ_PD_* = 3.5, the overall trend of DNA expansion with increasing *K_θ_* and stronger compaction at lower *λ_PP_* is preserved. In contrast, under moderate heterotypic interactions (*λ_PD_* = 0.5), DNA compaction is generally minimal, particularly at higher *K_θ_* values (very high *R_g_* values, comparable to the *R_g_* of a single DNA chain in the absence of proteins). Moreover, the ordering of compaction among the different *λ_PP_* values shifts, with *λ_PP_* = 3.0 showing more compaction than *λ_PP_* = 4.0, consistent with trends observed for DNA chain length.

**Figure 5:**
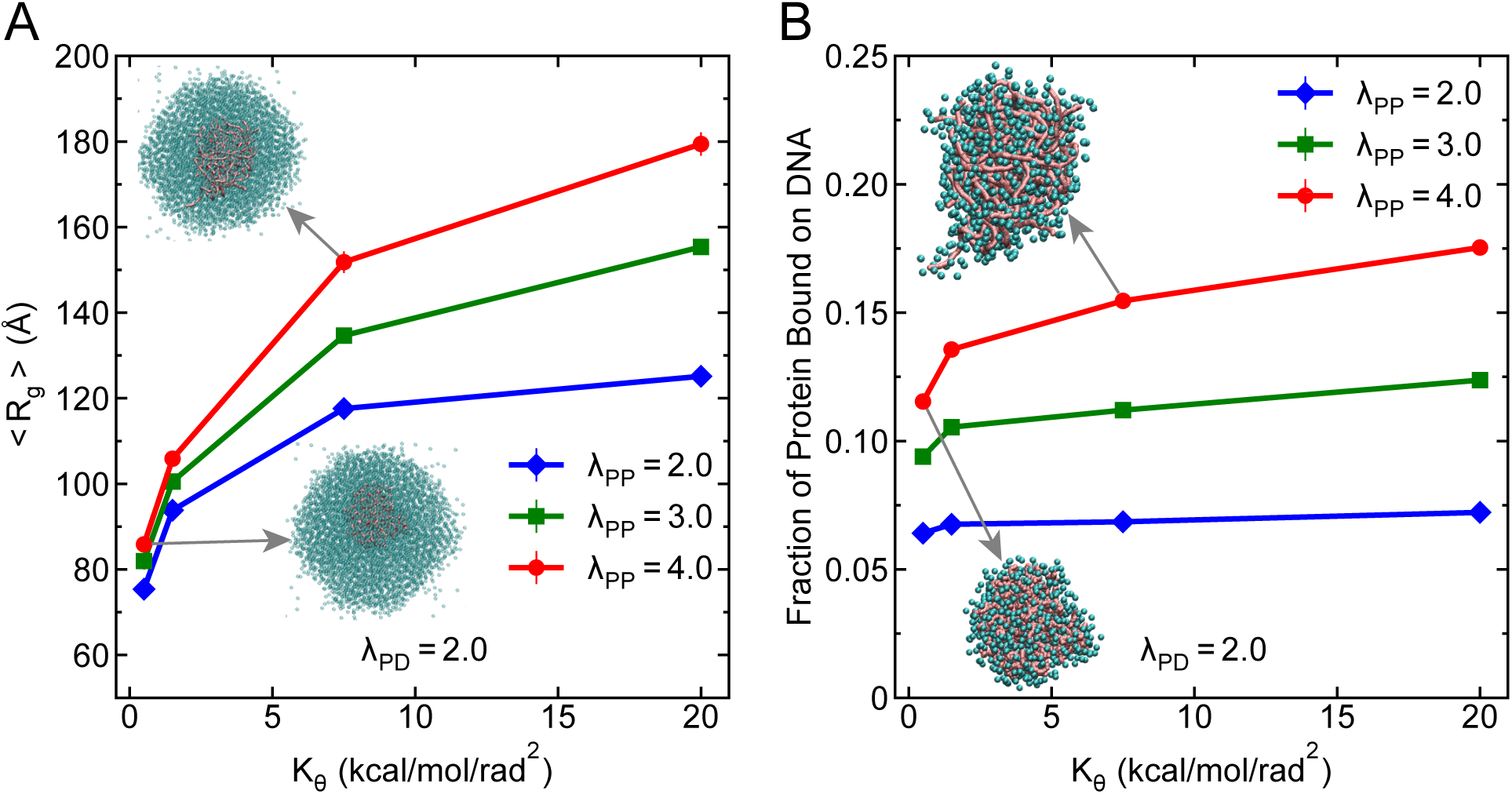
DNA flexibility affects the organization of proteins and DNA within the condensates. (A) Average *R_g_* of DNA as a function of bending rigidity (*K_θ_*) at a fixed heterotypic interaction strength *λ_PD_* = 2.0 for three different homotypic interaction strengths *λ_PP_* = 2.0, 3.0, and 4.0. Insets show representative structural organizatin of DNA chains within the protein-DNA condensate at two selected regidity values. (B) Fraction of proteins bound to DNA within the dense phase as a function of *K_θ_* for the same parameter sets as in panel A. Insets show the structural organization of proteins bound with DNA at two selected regidity values. DNA molecules are represented in pink color and protein molecules are represented in cyan color. Simulations used 250 bp long DNA chains (20 in total) and 5000 protein beads. In both panels, error bars represent standard errors obtained from block averaging and the associated error bars are smaller than the symbol size.

While comparing the protein bound to DNA for different DNA chain flexibility (Fig. 5B), we observed that protein fraction bound to DNA stays constant when the DNA acts as a scaffold to recruit protein and forms condensate (*λ_PP_* = 2.0). This observation can be related to the monolayer adsorption of FUS protein on ssDNA and dsDNA, where protein binding to DNA chain under lower protein concentration remains same independent of flexibility which is proposed as the mechanism for phase separation.^17^ Increasing homotypic interaction strength further leads to higher DNA-bound protein fractions for all *K_θ_* values (Fig. 5B), suggesting that enhanced homotypic interactions facilitate more favorable DNA accessibility within the condensate. The same qualitative trends are reproduced in simulations at moderate as well as at strong heterotypic strengths (Figs. S2B, D).

### Multiphasic Nature of Protein-DNA Condensates at Higher Heterotypic Interaction Strengths

While the above results provided the crucial insight into the role played by the DNA physical properties in organizing the condensate, we next explore how the mere presence of DNA influences the overall phase separation behavior of proteins. Recent studies have shown that the propensity of phase separation of proteins increases with incorporation of DNA molecules in condensates as indicated by the decrease in saturation concentrations of proteins.^22,27,32^ Thus, in order to understand how the phase behavior of proteins changes in the presence of DNA, we varied the interaction strength *λ_PP_* between proteins and calculated the coexistence concentrations of proteins in the dense and dilute phases (see Supporting Information for details). For this, we performed simulations under two different conditions: one with moderate protein–DNA interaction (*λ_PD_* = 0.5) and another with strong protein–DNA interaction (*λ_PD_* = 2.0) strength. In both cases, the system contained 5000 protein molecules and 20 DNA chains, each 250 base pairs long. As a reference, we also simulated a system with proteins only (without any DNA), see Fig. 1D. The results are shown in Fig. 6A. When the protein–DNA interaction is moderate (*λ_PD_* = 0.5), the phase diagram looks similar to that of the protein-only system. The dense and dilute phase concentrations converge to form a typical two-phase coexistence envelope (Fig. 1D), indicating no major change in phase behavior. However, when the protein–DNA interaction is strong (*λ_PD_* = 2.0), we observe a striking difference: the phase envelope does not converge (see also Fig. S3 for other DNA lengths). This non-converging behavior suggests that the nature of the condensate is fundamentally different under strong heterotypic interactions.

**Figure 6:**
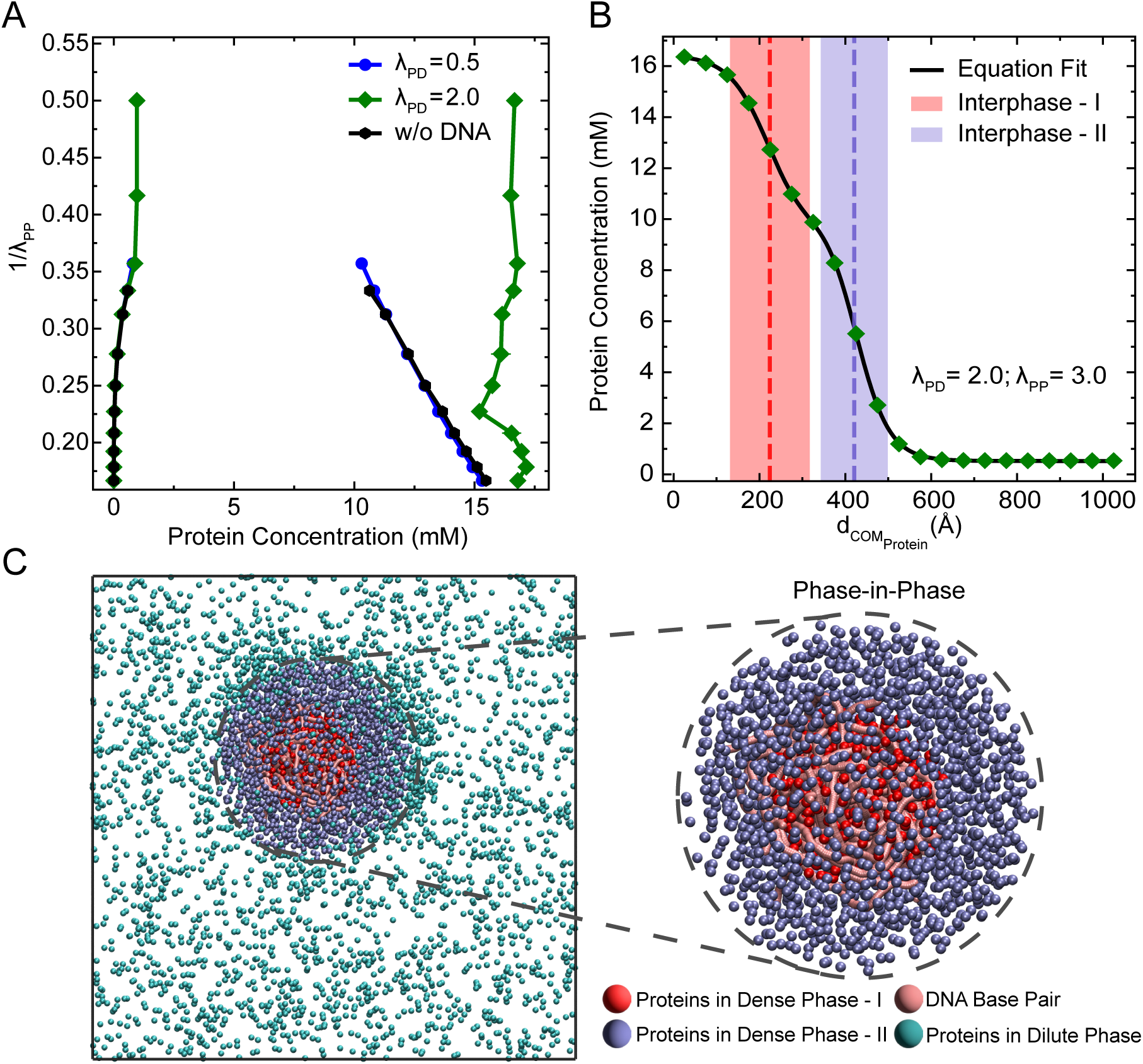
Multiphasic nature of protein-DNA condensates. (A) Phase diagrams showing protein concentrations in the dilute and dense phases as a function of protein–protein interaction strength (*λ_PP_*), for three systems: proteins only (black), proteins with DNA at moderate (*λ_PD_* = 0.5, blue) and strong (*λ_PD_* = 2.0, green) heterotypic interaction strength. (B) Radial density profile of proteins from the center of mass (COM) of proteins in the largest cluster at *λ_PD_* = 2.0 and *λ_PP_* = 3.0, showing two distinct protein populations. The profile is fitted with a double hyperbolic tangent function (solid black line), capturing the presence of an inner DNA-associated protein-rich region (dense phase-I) and an outer protein-rich region not in contact with DNA (dense phase-II). Red and blue dotted lines represent the location of the two interphases between dense phase-I, dense phase-II and between dense phase-II and dilute phase, respectively. (C) Representative simulation snapshot corresponding to panel B, illustrating the multiphasic organization. DNA-bound proteins (dense phase-I) are shown in red, unbound proteins (dense phase-II) in blue, DNA molecules in pink, and proteins in the dilute phase in cyan.

To understand this difference more clearly, we analyzed the spatial distribution of proteins within the condensate at *λ_PD_* = 2.0 and *λ_PP_* = 3.0, a condition corresponding to the nonconverging regime (see also Fig. S4 for other combinations of *λ_PD_* and *λ_PP_*). Specifically, we looked at how the protein density changes with distance from the center of mass (COM) of proteins in the largest cluster (see Supporting Information). As shown in Fig. 6B, the radial density profile reveals two distinct regions of proteins within the condensate, which can be clearly captured by fitting a double hyperbolic tangent function to the radial density distribution (see Supporting Information for more details). These two regions represent two types of protein populations within the same condensate. The first population consists of proteins that are bound to DNA and clustered around it, forming the inner phase (Dense phase-I) differentiated within condensate by the interphase-I, shown as the red-shaded region in Fig. 6B. The second population consists of proteins that are not bound to DNA and are distributed away from the DNA, forming the outer phase (Dense phase-II) distinguished from dilute phase by the interphase-II, shown as the blue-shaded region. A simulation snapshot in Fig. 6C further illustrates this structure, where red-colored proteins are DNA-bound (Dense phase-I), and blue-colored proteins are unbound (Dense phase-II). This observation indicates that the condensate is not uniform in composition, but rather contains multiple subphases, a phenomenon usually referred to as “multiphasic organization”.^60–62^ In other words, even though all proteins are chemically identical, their interactions with DNA create a spatial separation within the condensate, leading to a “phase-in-phase” architecture where different regions have different protein densities. Such architectures of multiphasic condensate have been reported in nucleolus, that has distinct immiscible phases^5,63^ and for the case of a ternary mixture of protein HP1, histone protein H1, and DNA where a layered organization was formed in the condensate which may help with DNA packaging within the condensate. ^32^

### Differential Protein Dynamics Within the Protein-DNA Condensate

As we observed the formation of distinct protein phases within a protein-DNA condensate for strong heterotypic interaction strengths, we hypothesized that these distinct protein phases might exhibit different dynamic behaviors. To evaluate the dynamics of proteins for multiple phases, the use of mean square displacement (MSD) of proteins for calculation of the diffusion coefficient (*D*) poses challenges, as it would average all the distinct modes of diffusion. To overcome this limitation, we employed an alternative method to compute *D* using the probability distribution of displacements, *P* (*r, t*), in the radial direction (*r*) as a function of lag time *t*.^64^ This method captures the presence of multiple diffusion behaviors by fitting the displacement distribution to a sum of Gaussian components. Each component is characterized by a distinct diffusion coefficient (*D_i_*) and its corresponding fractional population (*p_i_*), described by the expression:

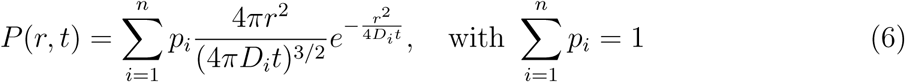

where *n* is the number of diffusivity (*D*) values that we increase for fitting the probability distribution until the fit achieves minimal residuals.

We find that it is necessary to include multiple diffusivity (*D*) values to fit the displacement distribution data, which indicates that there are different modes of diffusion for the proteins. For instance, we first applied this method to a system with weak heterotypic interaction strength (*λ_PD_* = 0.3) and strong homotypic interaction (*λ_PP_* = 3.0), under which DNA is not recruited in the condensate (see also Fig. 2). In Fig. S5A, the corresponding probability distribution is shown for different lag time *t* and found that we need three *D* values (*n* = 3) to best fit the displacement probability distribution, suggesting three separate modes of diffusion. Such best fitting is ensured from the residual, which is the difference between fitted values and the simulation data points, showing the minimal deviation across *r* (lower panel in Fig. S5A). We then checked these three *D* values obtained from the fitting, which remained constant across lag times, as shown in Fig. S5B, and likely correspond to: (i) fast-diffusing proteins in the dilute phase (*D*_1_), (ii) proteins at the interface between dense and dilute phases (*D*_2_), and (iii) slower-diffusing proteins deeply embedded in the condensed phase (*D*_3_).

Next, we analyzed this behavior for the multiphasic protein condensate formed under strong heterotypic (*λ_PD_* = 2.0) and homotypic (*λ_PP_* = 3.0) interaction strengths. Remarkably, in this case, we found that five distinct diffusivity modes (*n* = 5) were required for an optimal fit to accurately describe the displacement distributions (Fig. 7A and also Fig. S6), revealing a more complex dynamical landscape. These diffusivity components remained constant across lag times (Fig. 7B), indicating that each mode exhibits stable diffusivity across lag times and their averaged values are shown in Fig. 7C. Comparison with the DNA-excluded condensate clearly reveals two additional slow modes of diffusion (*D*_4_ and *D*_5_) in the DNA-recruited system (Fig. 7C). Based on the ordering of the diffusion coefficients (*D*_5_ *< D*_4_ *< D*_3_ *< D*_2_ *< D*_1_), we propose a possible interpretation of these five dynamic populations. The slowest diffusion component (*D*_5_) likely corresponds to proteins in the dense phase that are either immobilized by forming bridging contacts with DNA or are strongly bound to DNA. The next two modes (*D*_4_ and *D*_3_) may represent proteins bound on the DNA surface and those diffusing within the condensed phase but not in contact with DNA. The fourth diffusion mode (*D*_2_) corresponds to proteins at the interface of condensed and dilute phase and the fastest mode (*D*_1_) represent proteins in the dilute phase. These results thus emphasize that even within a spatially continuous condensate, proteins experience diverse microenvironments due to differential DNA interactions. Notably, such differential nature of protein dynamics has been experimentally observed in HP1*α* chromatin condensate where the proteins diffusivity varies with their distance from the chromatin in the condensate. ^65^ Experimentally, fluorescence recovery after photobleaching (FRAP) is the primary method used for determination of protein dynamics in condensates where a region of fluorescent proteins is photobleached and the time required for the recovery of fluorescence in this region is interpreted as a measure of protein dynamics.^66,67^ Our findings on differential protein dynamics within a condensate underscores the importance of selecting the appropriate region for studying the protein dynamics within condensate as different regions of proteins within condensate reflect different dynamic properties of proteins due to recruitment of DNA.

**Figure 7:**
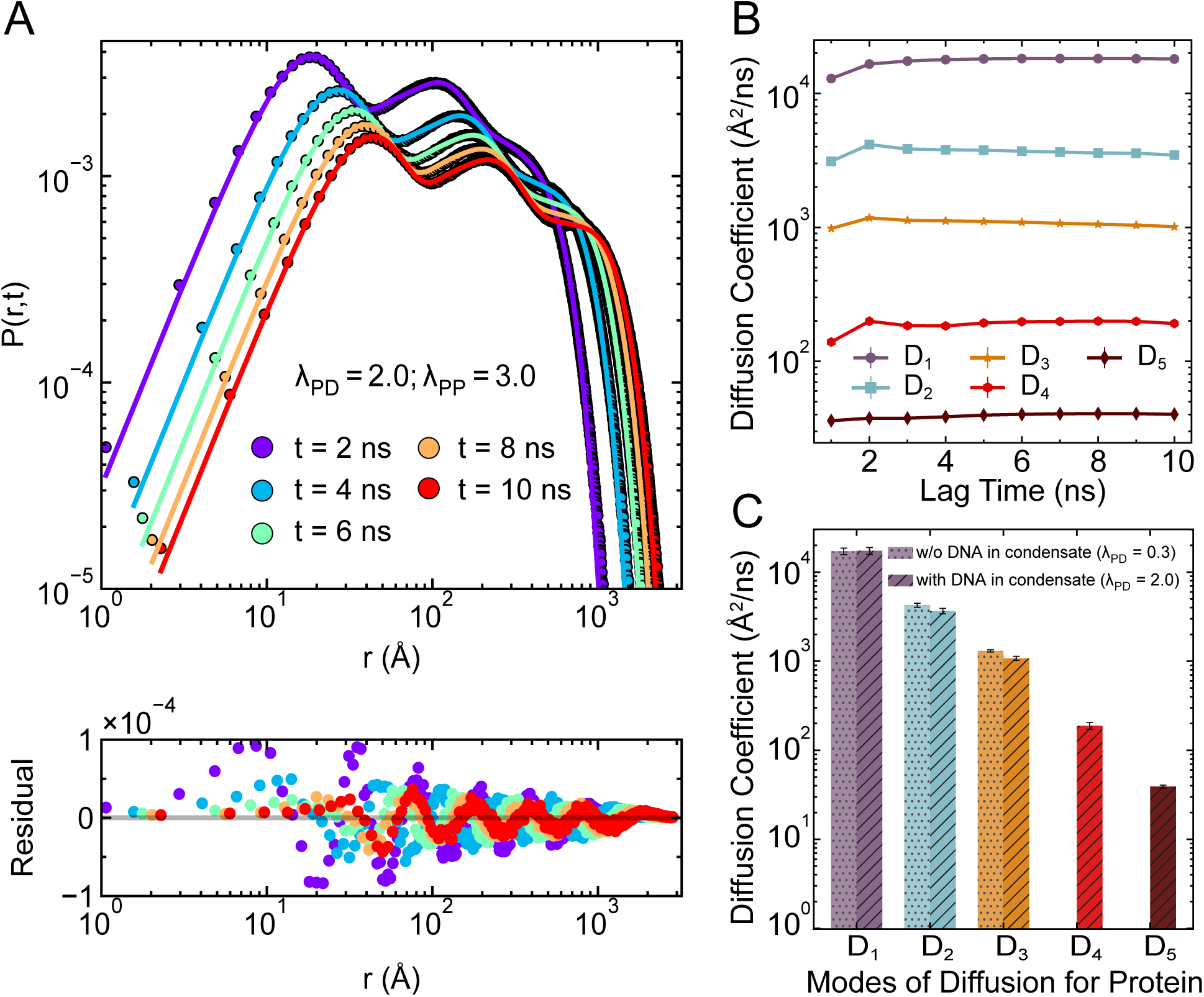
Differential dynamics of protein within protein-DNA condensates. (A) Radial displacement probability distributions of proteins, *P* (*r, t*), for different lag times *t*, under strong heterotypic (*λ_PD_* = 2.0) and homotypic (*λ_PP_* = 3.0) interaction strengths. The points denote histogram data from simulations, while solid lines show the best-fit using a sum of five Gaussian components (Eq. 6). The difference between the Gaussian fitted lines and the simulation data points are represented as residuals from the data fitting for each lag time *t* (lower panel), confirming the quality of fit (see also Fig. S6). (B) Diffusion coefficients (*D_i_*) corresponding to the five dynamic modes extracted from the Gaussian fits in (A), shown as a function of lag time. The plateau behavior indicates that each mode exhibits stable diffusivity across lag times. (C) Comparison of the average diffusion coefficients for different dynamic modes between the DNA-recruited condensate (*λ_PD_* = 2.0, hatched bars) and DNA-excluded condensate (*λ_PD_* = 0.3, dotted bars). The DNA-recruited case reveals two additional slow diffusion modes (*D*_4_ and *D*_5_), highlighting the emergence of new dynamic populations due to DNA localization within the condensate.

## Summary and Conclusions

In this study, we employed a minimalist coarse-grained model to investigate how heterotypic protein-DNA and homotypic protein-protein interactions, along with DNA physical properties such as chain length and flexibility, influence the formation, structure, and dynamics of protein-DNA condensates. Our simulations revealed that while homotypic interactions are sufficient to drive protein condensation, heterotypic interactions with DNA greatly influence the phase behavior and condensate formation. We found that DNA acts as a scaffold for recruiting proteins when heterotypic interactions are stronger, leading to the compaction of DNA and formation of condensate. We further demonstrated that longer DNA chains and more flexible DNA molecules undergo higher degree of compaction and promote the formation of distinct structural morphologies within condensates, including multiphasic organizations. These structural features are regulated by the strength of homotypic and heterotypic interactions.

Importantly, our results highlighted the existence of distinct protein microenvironments within the condensate, with protein populations exhibiting different binding modes, ranging from strongly DNA-bound to completely unbound and freely diffusing in the dilute phase. These diverse structural states lead to markedly different diffusion behaviors. By analyzing the distributions of radial displacements of proteins, we uncovered up to five distinct diffusion modes, with the slowest corresponding to strongly DNA-bound proteins and the fastest to freely diffusing ones. Such findings resonate with experimental observations of spatially dependent protein dynamics in chromatin-associated condensates and provide a physical basis for interpreting heterogeneous fluorescence recovery signals seen in FRAP experiments.

Our work demonstrates that DNA is not merely a passive scaffold but an active modulator of condensate architecture and dynamics, capable of recruiting proteins, altering their spatial organization, and modulating their mobility. This mechanistic understanding is crucial for interpreting the functional consequences of biomolecular condensates *in vivo*, particularly in chromatin organization, transcriptional regulation, and other DNA-templated processes.

While our CG simulation results present a plausible microscopic picture of the formation of protein-DNA condensates, it is important to discuss its limitations. The proteins and DNA are represented using simplified models that lack sequence specificity and do not capture detailed structural features such as helical twisting of DNA or the multi-domain architecture of real proteins. Furthermore, the model neglects explicit solvent and electrostatic effects, which may play important roles in determining the phase behavior and internal structuring of real biomolecular condensates. Moreover, our model also does not include specific interactions between proteins or between proteins and DNA such as multivalent domain-mediated contacts, DNA-binding motifs, or charge patterning effects, which are known to significantly influence condensate composition, selectivity, and material properties.^35,68,69^ Nevertheless, despite these simplifications, our model successfully recapitulates key qualitative features observed experimentally and provides a foundational framework for understanding the interplay between molecular properties and phase organization in protein-DNA condensates. This work lays the groundwork for future studies incorporating more detailed molecular representations and sequence-specific interactions to bridge the gap between minimal models and biological complexity.

## Supporting Information Available

Calculation of DNA persistence length; calculation of radius of gyration (*R_g_*) of DNA chains; analysis of radial density profile of proteins in single and multiphasic condensates; normalized *R_g_*of DNA molecules and fraction of proteins bound to DNA as a function of DNA length for moderate (*λ_PD_* = 0.5) and strong (*λ_PD_*= 3.5) heterotypic interaction strengths with three different homotypic interaction strengths; average *R_g_* of DNA of length 250 bp and fraction of proteins bound to DNA as a function of bending rigidity *K_θ_*for moderate (*λ_PD_* = 0.5) and strong (*λ_PD_*= 3.5) heterotypic interaction strengths with three different homotypic interaction strengths; phase diagram of protein concentrations in the dilute and dense phases as a function of *λ_PP_* for three different DNA lengths at strong *λ_PD_* = 2.0; radial density profile of proteins from the center of mass of proteins in the largest cluster at *λ_PD_* = 2.5, *λ_PP_* = 3.6 and *λ_PD_* = 2.5, *λ_PP_* = 4.0; radial displacement probability distributions of proteins for different lag times at *λ_PD_* = 0.3, *λ_PP_* = 3.0 and diffusion coefficients corresponding to the three dynamic modes of proteins as a function of lag time; residual analysis for Gaussian decomposition of protein displacement distribution in multiphase condensates.

## Author Contributions

A.S.T and J.M. designed research; A.S.T performed research; A.S.T, A.M., J.W., Y.C.K and J.M. analyzed data; and A.S.T, A.M., and J.M. wrote the paper.

## Acknowledgement

The work presented in this article has been supported by NIH/NIGMS R35GM153388.

Y.C.K. is supported by the Office of Naval Research through the U.S. Naval Research Laboratory base program. We acknowledge the Texas A & M High Performance Research Computing (HPRC) for providing computational resources required for performing this work.

## Data and Software Availability

All data are included in the manuscript and/or supporting information.

## References

(1) Shin, Y.; Brangwynne, C. P. Liquid phase condensation in cell physiology and disease. Science 2017, 357, eaaf4382.

(2) Banani, S. F.; Lee, H. O.; Hyman, A. A.; Rosen, M. K. Biomolecular condensates: organizers of cellular biochemistry. Nature reviews Molecular cell biology 2017, 18, 285–298.

(3) Mao, Y. S.; Zhang, B.; Spector, D. L. Biogenesis and function of nuclear bodies. Trends in Genetics 2011, 27, 295–306.

(4) Decker, C. J.; Parker, R. P-bodies and stress granules: possible roles in the control of translation and mRNA degradation. Cold Spring Harbor perspectives in biology 2012, 4, a012286.

(5) Feric, M.; Vaidya, N.; Harmon, T. S.; Mitrea, D. M.; Zhu, L.; Richardson, T. M.; Kriwacki, R. W.; Pappu, R. V.; Brangwynne, C. P. Coexisting liquid phases underlie nucleolar subcompartments. Cell 2016, 165, 1686–1697.

(6) Feric, M.; Demarest, T. G.; Tian, J.; Croteau, D. L.; Bohr, V. A.; Misteli, T. Self-assembly of multi-component mitochondrial nucleoids via phase separation. The EMBO journal 2021, 40, e107165.

(7) Hyman, A. A.; Weber, C. A.; Jülicher, F. Liquid-liquid phase separation in biology. Annual review of cell and developmental biology 2014, 30, 39–58.

(8) Li, P.; Banjade, S.; Cheng, H.-C.; Kim, S.; Chen, B.; Guo, L.; Llaguno, M.; Hollingsworth, J. V.; King, D. S.; Banani, S. F.; Russo, P. S.; Jiang, Q.-X.; Nixon, B. T.; Rosen, M. K. Phase transitions in the assembly of multivalent signalling proteins. Nature 2012, 483, 336–340.

(9) King, J. T.; Shakya, A. Phase separation of DNA: From past to present. Biophysical journal 2021, 120, 1139–1149.

(10) Dignon, G. L.; Best, R. B.; Mittal, J. Biomolecular phase separation: from molecular driving forces to macroscopic properties. Annual review of physical chemistry 2020, 71, 53–75.

(11) Murthy, A. C.; Tang, W. S.; Jovic, N.; Janke, A. M.; Seo, D. H.; Perdikari, T. M.; Mittal, J.; Fawzi, N. L. Molecular interactions contributing to FUS SYGQ LC-RGG phase separation and co-partitioning with RNA polymerase II heptads. Nature structural & molecular biology 2021, 28, 923–935.

(12) Mondal, A.; Bhattacherjee, A. Searching target sites on DNA by proteins: Role of DNA dynamics under confinement. Nucleic Acids Research 2015, 43, 9176–9186.

(13) Mohanty, P.; Kapoor, U.; Sundaravadivelu Devarajan, D.; Phan, T. M.; Rizuan, A.; Mittal, J. Principles governing the phase separation of multidomain proteins. Biochemistry 2022, 61, 2443–2455.

(14) Rekhi, S.; Garcia, C. G.; Barai, M.; Rizuan, A.; Schuster, B. S.; Kiick, K. L.; Mittal, J. Expanding the molecular language of protein liquid–liquid phase separation. Nature chemistry 2024, 16, 1113–1124.

(15) Larson, A. G.; Elnatan, D.; Keenen, M. M.; Trnka, M. J.; Johnston, J. B.; Burlingame, A. L.; Agard, D. A.; Redding, S.; Narlikar, G. J. Liquid droplet formation by HP1*α* suggests a role for phase separation in heterochromatin. Nature 2017, 547, 236–240.

(16) Laghmach, R.; Di Pierro, M.; Potoyan, D. A. The interplay of chromatin phase separation and lamina interactions in nuclear organization. Biophysical Journal 2021, 120, 5005–5017.

(17) Renger, R.; Morin, J. A.; Lemaitre, R.; Ruer-Gruss, M.; Jülicher, F.; Hermann, A.; Grill, S. W. Co-condensation of proteins with single- and double-stranded DNA. Proceedings of the National Academy of Sciences 2022, 119, e2107871119.

(18) Shrinivas, K.; Sabari, B. R.; Coffey, E. L.; Klein, I. A.; Boija, A.; Zamudio, A. V.; Schuijers, J.; Hannett, N. M.; Sharp, P. A.; Young, R. A.; Chakraborty, A. K. Enhancer features that drive formation of transcriptional condensates. Molecular cell 2019, 75, 549–561.

(19) Qamar, S.; Wang, G.; Randle, S. J.; Ruggeri, F. S.; Varela, J. A.; Lin, J. Q.; Phillips, E. C.; Miyashita, A.; Williams, D.; Ströhl, F.; others FUS phase separation is modulated by a molecular chaperone and methylation of arginine cation-*π* interactions. Cell 2018, 173, 720–734.

(20) Gibson, B. A.; Doolittle, L. K.; Schneider, M. W.; Jensen, L. E.; Gamarra, N.; Henry, L.; Gerlich, D. W.; Redding, S.; Rosen, M. K. Organization of chromatin by intrinsic and regulated phase separation. Cell 2019, 179, 470–484.

(21) Brackley, C. A.; Liebchen, B.; Michieletto, D.; Mouvet, F.; Cook, P. R.; Marenduzzo, D. Ephemeral protein binding to DNA shapes stable nuclear bodies and chromatin domains. Biophysical journal 2017, 112, 1085–1093.

(22) Phan, T. M.; Kim, Y. C.; Debelouchina, G. T.; Mittal, J. Interplay between charge distribution and DNA in shaping HP1 paralog phase separation and localization. Elife 2024, 12, RP90820.

(23) Her, C.; Phan, T. M.; Jovic, N.; Kapoor, U.; Ackermann, B. E.; Rizuan, A.; Kim, Y. C.; Mittal, J.; Debelouchina, G. T. Molecular interactions underlying the phase separation of HP1*α*: role of phosphorylation, ligand and nucleic acid binding. Nucleic Acids Research 2022, 50, 12702–12722.

(24) Quail, T.; Golfier, S.; Elsner, M.; Ishihara, K.; Murugesan, V.; Renger, R.; Jülicher, F.; Brugués, J. Force generation by protein-DNA co-condensation. Nature Physics 2021, 17, 1007–1012.

(25) Monahan, Z.; Ryan, V. H.; Janke, A. M.; Burke, K. A.; Rhoads, S. N.; Zerze, G. H.; O’Meally, R.; Dignon, G. L.; Conicella, A. E.; Zheng, W.; others Phosphorylation of the FUS low-complexity domain disrupts phase separation, aggregation, and toxicity. The EMBO journal 2017, 36, 2951–2967.

(26) Hazra, M. K.; Levy, Y. Cross-talk of cation-*π* interactions with electrostatic and aromatic interactions: A salt-dependent trade-off in biomolecular condensates. The journal of physical chemistry letters 2023, 14, 8460–8469.

(27) Keenen, M. M.; Brown, D.; Brennan, L. D.; Renger, R.; Khoo, H.; Carlson, C. R.; Huang, B.; Grill, S. W.; Narlikar, G. J.; Redding, S. HP1 proteins compact DNA into mechanically and positionally stable phase separated domains. elife 2021, 10, e64563.

(28) Ryu, J.-K.; Bouchoux, C.; Liu, H. W.; Kim, E.; Minamino, M.; de Groot, R.; Katan, A. J.; Bonato, A.; Marenduzzo, D.; Michieletto, D.; Uhlmann, F.; Dekker, C. Bridging-induced phase separation induced by cohesin SMC protein complexes. Science advances 2021, 7, eabe5905.

(29) Shakya, A.; King, J. T. DNA local-flexibility-dependent assembly of phase-separated liquid droplets. Biophysical journal 2018, 115, 1840–1847.

(30) Kapoor, U.; Kim, Y. C.; Mittal, J. Coarse-grained models to study protein–DNA interactions and liquid–liquid phase separation. Journal of Chemical Theory and Computation 2023, 20, 1717–1731.

(31) Yasuda, I.; von Bülow, S.; Tesei, G.; Yamamoto, E.; Yasuoka, K.; Lindorff-Larsen, K. Coarse-grained model of disordered RNA for simulations of biomolecular condensates. Journal of chemical theory and computation 2025, 21, 2766–2779.

(32) Latham, A. P.; Zhang, B. On the stability and layered organization of protein-DNA condensates. Biophysical Journal 2022, 121, 1727–1737.

(33) Ancona, M.; Brackley, C. A. Simulating the chromatin-mediated phase separation of model proteins with multiple domains. Biophysical Journal 2022, 121, 2600–2612.

(34) Dignon, G. L.; Zheng, W.; Kim, Y. C.; Best, R. B.; Mittal, J. Sequence determinants of protein phase behavior from a coarse-grained model. PLoS computational biology 2018, 14, e1005941.

(35) Sundaravadivelu Devarajan, D.; Wang, J.; Sza-la-Mendyk, B.; Rekhi, S.; Nikoubashman, A.; Kim, Y. C.; Mittal, J. Sequence-dependent material properties of biomolecular condensates and their relation to dilute phase conformations. Nature Communications 2024, 15, 1912.

(36) Miller, C. M.; Kim, Y. C.; Mittal, J. Protein composition determines the effect of crowding on the properties of disordered proteins. Biophysical journal 2016, 111, 28– 37.

(37) Welles, R. M.; Sojitra, K. A.; Garabedian, M. V.; Xia, B.; Wang, W.; Guan, M.; Regy, R. M.; Gallagher, E. R.; Hammer, D. A.; Mittal, J.; others Determinants that enable disordered protein assembly into discrete condensed phases. Nature Chemistry 2024, 16, 1062–1072.

(38) Statt, A.; Casademunt, H.; Brangwynne, C. P.; Panagiotopoulos, A. Z. Model for disordered proteins with strongly sequence-dependent liquid phase behavior. The Journal of chemical physics 2020, 152, 075101.

(39) Rana, U.; Brangwynne, C. P.; Panagiotopoulos, A. Z. Phase separation vs aggregation behavior for model disordered proteins. The Journal of chemical physics 2021, 155, 125101.

(40) Brackley, C. A.; Taylor, S.; Papantonis, A.; Cook, P. R.; Marenduzzo, D. Nonspecific bridging-induced attraction drives clustering of DNA-binding proteins and genome organization. Proceedings of the National Academy of Sciences 2013, 110, E3605–E3611.

(41) Rubio-Cosials, A.; Sydow, J. F.; Jiménez-Menéndez, N.; Fernández-Millán, P.; Montoya, J.; Jacobs, H. T.; Coll, M.; Bernadó, P.; Solà, M. Human mitochondrial transcription factor A induces a U-turn structure in the light strand promoter. Nature structural & molecular biology 2011, 18, 1281–1289.

(42) Manning, G. S. The persistence length of DNA is reached from the persistence length of its null isomer through an internal electrostatic stretching force. Biophysical journal 2006, 91, 3607–3616.

(43) Mitchell, J. S.; Glowacki, J.; Grandchamp, A. E.; Manning, R. S.; Maddocks, J. H. Sequence-dependent persistence lengths of DNA. Journal of chemical theory and computation 2017, 13, 1539–1555.

(44) Ashbaugh, H. S.; Hatch, H. W. Natively unfolded protein stability as a coil-to-globule transition in charge/hydropathy space. Journal of the American Chemical Society 2008, 130, 9536–9542.

(45) Anderson, J. A.; Lorenz, C. D.; Travesset, A. General purpose molecular dynamics simulations fully implemented on graphics processing units. Journal of computational physics 2008, 227, 5342–5359.

(46) Anderson, J. A.; Glaser, J.; Glotzer, S. C. HOOMD-blue: A Python package for highperformance molecular dynamics and hard particle Monte Carlo simulations. Computational Materials Science 2020, 173, 109363.

(47) Ramasubramani, V.; Dice, B. D.; Harper, E. S.; Spellings, M. P.; Anderson, J. A.; Glotzer, S. C. freud: A software suite for high throughput analysis of particle simulation data. Computer Physics Communications 2020, 254, 107275.

(48) Patel, A.; Lee, H. O.; Jawerth, L.; Maharana, S.; Jahnel, M.; Hein, M. Y.; Stoynov, S.; Mahamid, J.; Saha, S.; Franzmann, T. M.; others A liquid-to-solid phase transition of the ALS protein FUS accelerated by disease mutation. Cell 2015, 162, 1066–1077.

(49) Murthy, A. C.; Dignon, G. L.; Kan, Y.; Zerze, G. H.; Parekh, S. H.; Mittal, J.; Fawzi, N. L. Molecular interactions underlying liquid-liquid phase separation of the FUS low-complexity domain. Nature structural & molecular biology 2019, 26, 637–648.

(50) Nott, T. J.; Petsalaki, E.; Farber, P.; Jervis, D.; Fussner, E.; Plochowietz, A.; Craggs, T. D.; Bazett-Jones, D. P.; Pawson, T.; Forman-Kay, J. D.; Baldwin, A. J. Phase transition of a disordered nuage protein generates environmentally responsive membraneless organelles. Molecular cell 2015, 57, 936–947.

(51) Stortz, M.; Presman, D. M.; Levi, V. Transcriptional condensates: a blessing or a curse for gene regulation? Communications Biology 2024, 7, 187.

(52) Brackley, C. A.; Marenduzzo, D. Bridging-induced microphase separation: photobleaching experiments, chromatin domains and the need for active reactions. Briefings in functional genomics 2020, 19, 111–118.

(53) Shahu, S.; Vtyurina, N.; Das, M.; Meyer, A. S.; Ganji, M.; Abbondanzieri, E. A. Bridging DNA contacts allow Dps from E. coli to condense DNA. Nucleic Acids Research 2024, 52, 4456–4465.

(54) Morin, J. A.; Wittmann, S.; Choubey, S.; Klosin, A.; Golfier, S.; Hyman, A. A.; Jülicher, F.; Grill, S. W. Sequence-dependent surface condensation of a pioneer transcription factor on DNA. Nature physics 2022, 18, 271–276.

(55) Ahlawat, V.; Dhiman, A.; Mudiyanselage, H. E.; Zhou, H.-X. Protamine-mediated tangles produce extreme deoxyribonucleic acid compaction. Journal of the American Chemical Society 2024, 146, 30668–30677.

(56) Chhetri, K. B.; Jang, Y. H.; Lansac, Y.; Maiti, P. K. Effect of phosphorylation of protamine-like cationic peptide on the binding affinity to DNA. Biophysical Journal 2022, 121, 4830–4839.

(57) Kim, Y. J.; Lee Jr, M.; Lee, Y.-T.; Jing, J.; Sanders, J. T.; Botten, G. A.; He, L.; Lyu, J.; Zhang, Y.; Mettlen, M.; Ly, P.; Zhou, Y.; Xu, J. Light-activated macromolecular phase separation modulates transcription by reconfiguring chromatin interactions. Science Advances 2023, 9, eadg1123.

(58) André, A. A.; Spruijt, E. Rigidity rules in DNA droplets: Nucleic acid flexibility affects model membraneless organelles. Biophysical Journal 2018, 115, 1837–1839.

(59) Shakya, A.; Girard, M.; King, J. T.; Olvera de la Cruz, M. Role of chain flexibility in asymmetric polyelectrolyte complexation in salt solutions. Macromolecules 2020, 53, 1258–1269.

(60) Kelley, F. M.; Favetta, B.; Regy, R. M.; Mittal, J.; Schuster, B. S. Amphiphilic proteins coassemble into multiphasic condensates and act as biomolecular surfactants. Proceedings of the National Academy of Sciences 2021, 118, e2109967118.

(61) Pandey, V.; Hosokawa, T.; Hayashi, Y.; Urakubo, H. Multiphasic protein condensation governed by shape and valency. Cell Reports 2025, 44, 115504.

(62) Rana, U.; Xu, K.; Narayanan, A.; Walls, M. T.; Panagiotopoulos, A. Z.; Avalos, J. L.; Brangwynne, C. P. Asymmetric oligomerization state and sequence patterning can tune multiphase condensate miscibility. Nature chemistry 2024, 16, 1073–1082.

(63) Lafontaine, D. L.; Riback, J. A.; Bascetin, R.; Brangwynne, C. P. The nucleolus as a multiphase liquid condensate. Nature reviews Molecular cell biology 2021, 22, 165–182.

(64) Zheng, W.; Dignon, G. L.; Jovic, N.; Xu, X.; Regy, R. M.; Fawzi, N. L.; Kim, Y. C.; Best, R. B.; Mittal, J. Molecular details of protein condensates probed by microsecond long atomistic simulations. The Journal of Physical Chemistry B 2020, 124, 11671– 11679.

(65) Strom, A. R.; Emelyanov, A. V.; Mir, M.; Fyodorov, D. V.; Darzacq, X.; Karpen, G. H. Phase separation drives heterochromatin domain formation. Nature 2017, 547, 241– 245.

(66) Taylor, N. O.; Wei, M.-T.; Stone, H. A.; Brangwynne, C. P. Quantifying dynamics in phase-separated condensates using fluorescence recovery after photobleaching. Biophysical journal 2019, 117, 1285–1300.

(67) Soranno, A. The trap in the FRAP: a cautionary tale about transport measurements in biomolecular condensates. Biophysical journal 2019, 117, 2041–2042.

(68) Devarajan, D. S.; Rekhi, S.; Nikoubashman, A.; Kim, Y. C.; Howard, M. P.; Mittal, J. Effect of charge distribution on the dynamics of polyampholytic disordered proteins. Macromolecules 2022, 55, 8987–8997.

(69) Maurici, N.; Phan, T. M.; Henty-Ridilla, J. L.; Kim, Y. C.; Mittal, J.; Bah, A. Uncovering the molecular interactions underlying MBD2 and MBD3 phase separation. The Journal of Physical Chemistry B 2025,

