## Supplementary Information for "Multiphasic Organization and Differential Dynamics of Proteins Within Protein-DNA Biomolecular Condensates"

In this supporting information, we present supporting text and additional figures that were referenced in the main text.

### Calculation of DNA Persistence Length

The persistence length ($l_p$) of DNA chains is calculated using the polymer analysis library from the MDAnalysis[1] python package. This approach relies on evaluating the orientational correlations between bond vectors along the polymer. The autocorrelation function $C(n)$ between bond vectors separated by $n$ bonds is computed as:

$$C(n) = \langle \cos\theta_{i,i+n} \rangle = \langle \mathbf{a_i} \cdot \mathbf{a_{i+n}} \rangle \tag{S1}$$

where $\mathbf{a_i}$ and $\mathbf{a_{i+n}}$ are unit vectors along bonds $i$ and $i+n$, respectively, and the angular bracket denotes an ensemble average over all pairs separated by $n$ bonds.

To extract the persistence length, the computed autocorrelation function is fit to the theoretical exponential decay:

$$C(n) \approx \exp\left(-\frac{nl_b}{l_p}\right) \tag{S2}$$

where $l_b = 5.5$ Å is the average bond length between adjacent DNA beads. By fitting Eq. (S2) to the data obtained from Eq. (S1), the persistence length $l_p$ of the DNA chain is estimated.

### Radius of Gyration Calculations for DNA Molecules

We computed the radius of gyration ($R_g$) of DNA chains by calculating the gyration tensor as[2] :

$$\mathbf{G} = \frac{1}{M}\sum_{i=1}^{N} m_i\,\Delta\mathbf{r}_i\,\Delta\mathbf{r}_i \tag{S3}$$

where $\Delta\mathbf{r}_i$ is the vector to $i^{th}$ monomer from the chain's center of mass, $m_i$ is the mass of the $i^{th}$ monomer, $M$ is the total mass of the chain and $N$ is the total number of monomer. From this gyration tensor, we calculate the radius of gyration of the DNA chains as the square root of the trace of the gyration tensor $R_g = \langle \mathrm{tr}\,\mathbf{G} \rangle^{1/2}$.

### Radial Density Profile Analysis of Proteins in Single and Multiphasic Condensates

For calculating the density profile of proteins across the dense and dilute phase, we calculate the largest cluster of proteins using Freud[3] Python library (version 2.12.1) with a distance criterion where protein molecules within 1.5 $\sigma_P$ are considered part of the same cluster, where $\sigma_P = 50$ Å is the diameter of the protein bead. The center of mass (COM) of the proteins in this largest cluster is calculated, and that position is regarded as the origin coordinate. From the COM, the distance of all protein beads is calculated, and the distribution of this distance is binned with a width of $\sigma_P$ to generate the radial density distribution of proteins. This radial density distribution is then fitted to a hyperbolic tangent function,[4] shown in Eq. (S4) to determine the dense phase concentration, the dilute phase concentration, and the location and width of the interface.

$$C(r) = \frac{1}{2}\big(C_{dense} + C_{dilute}\big) - \frac{1}{2}\big(C_{dense} - C_{dilute}\big)tanh\left(\frac{2(r - b_{int})}{c_{int}}\right) \tag{S4}$$

where $C(r)$ is the concentration of protein, $C_{dense}$ is the dense phase protein concentration, $C_{dilute}$ is the dilute phase protein concentration, $b_{int}$ is the location of the interface and $c_{int}$ is the width of interface.

For evaluating the protein concentration in case of multiphasic condensate formation, the radial density distribution is calculated as mentioned above but this distribution is then fitted to a combination of two hyperbolic tangent functions, shown in Eq. (S5) to determine the concentration of both the dense phases, the dilute phase, the width and the location of the two interfaces:

$$C(r) = A(r) \times switch(r) + B(r) \times \big(1 - switch(r)\big) \tag{S5}$$

where

$$A(r) = \frac{1}{2}\big(C_{dense,I} + C_{dense,II}\big) - \frac{1}{2}\big(C_{dense,I} - C_{dense,II}\big)tanh\left(\frac{2(r - b_{int,I})}{c_{int,I}}\right) \tag{S6}$$

$$B(r) = \frac{1}{2}\big(C_{dense,II} + C_{dilute}\big) - \frac{1}{2}\big(C_{dense,II} - C_{dilute}\big)tanh\left(\frac{2(r - b_{int,II})}{c_{int,II}}\right) \tag{S7}$$

and

$$switch(r) = \frac{1}{2}\left(1 - tanh\left(\frac{2(r - \frac{b_{int,I}+b_{int,II}}{2})}{b_{int,II} - b_{int,I}}\right)\right) \tag{S8}$$

Here, $C_{dense,I}$ is the first dense phase protein concentration, $C_{dense,II}$ is the second dense phase protein concentration, $C_{dilute}$ is the dilute phase protein concentration, $b_{int,I}$ and $b_{int,II}$ are the location of the first and second interfaces, $c_{int,I}$ and $c_{int,II}$ are the width of both interfaces.

#### Supplementary Figures

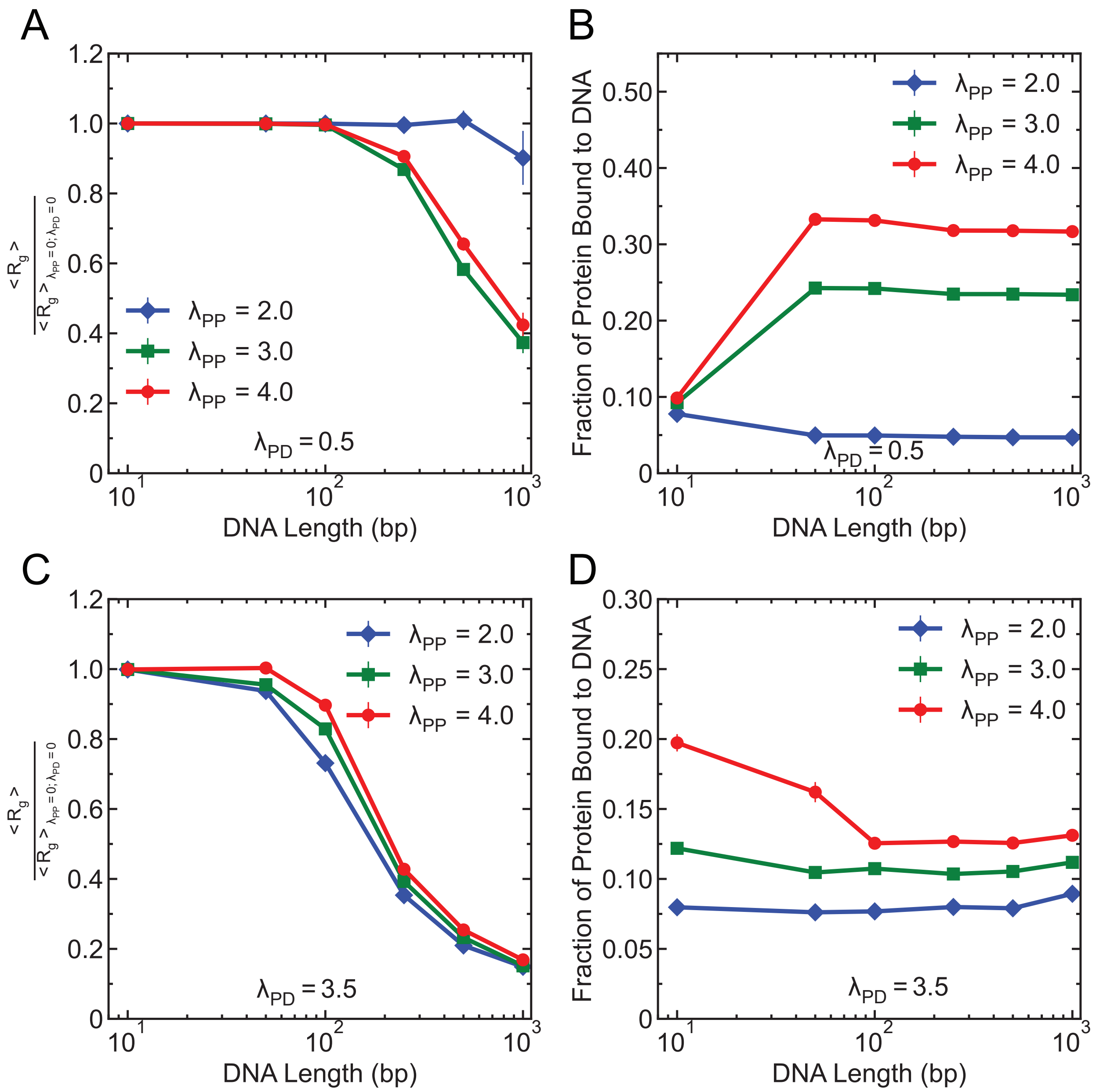


Figure S1: (A) Normalized radius of gyration ($R_g$) of DNA molecules as a function of DNA length for $\lambda_{PD} = 0.5$ with three different homotypic interaction strengths $\lambda_{PP} = 2.0, 3.0, 4.0$. $R_g$ is normalized by the $R_g$ of corresponding DNA chains in the absence of attractive interactions ($\lambda_{PP} = \lambda_{PD} = 0$). (B) Fraction of proteins bound to DNA as a function of DNA length under the same conditions as in panel (A). (C) Normalized $R_g$ of DNA as a function of DNA chain length for $\lambda_{PD} = 3.5$ with three different homotypic strengths $\lambda_{PP}$. (D) Fraction of proteins bound to DNA as a function of DNA length under the same conditions as in panel (C).

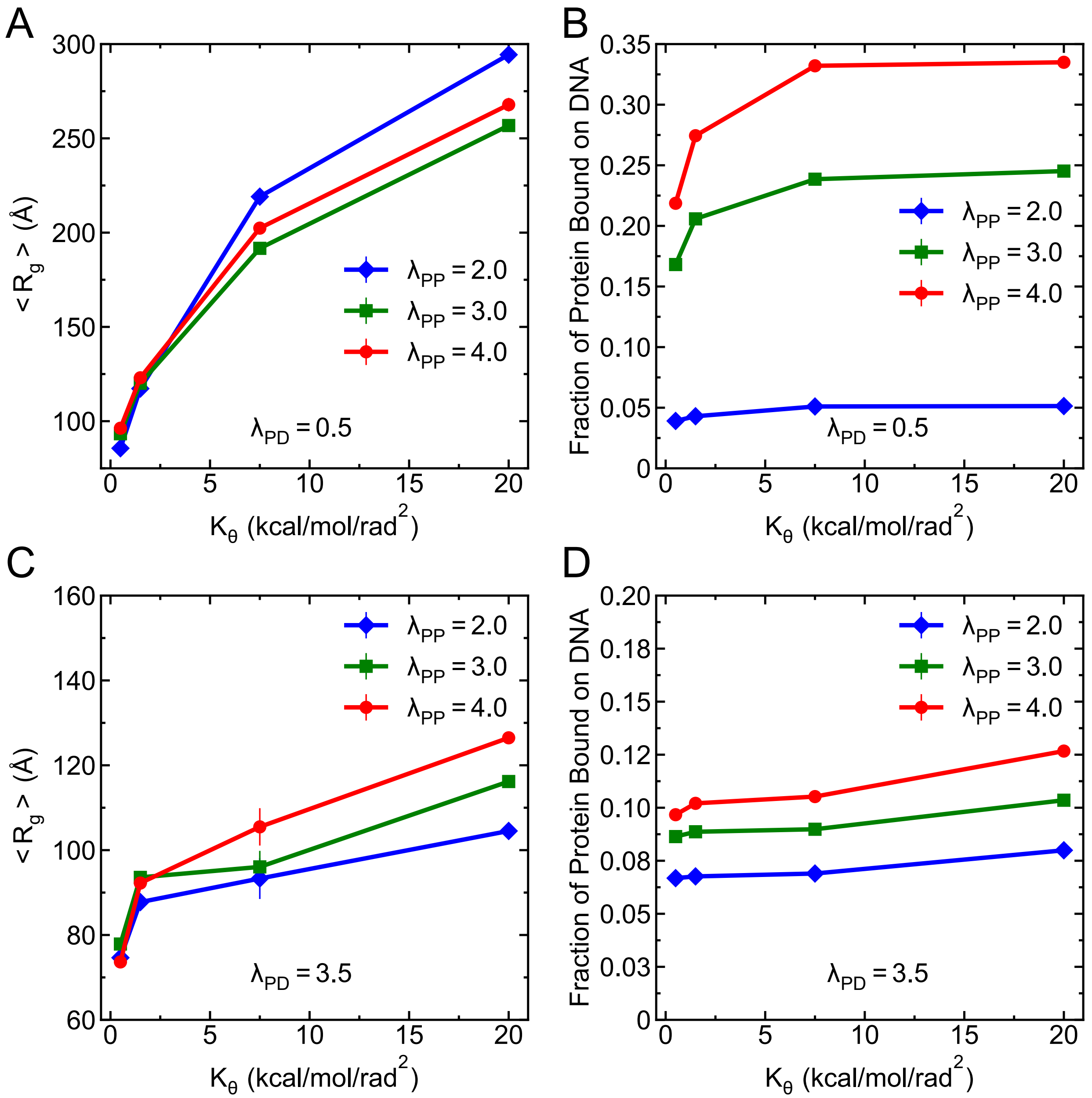


Figure S2: (A) Average $R_g$ of DNA of length 250 bp as a function of bending rigidity ($K_\theta$) for $\lambda_{PD} = 0.5$ with three different homotypic interaction strengths $\lambda_{PP} = 2.0, 3.0, 4.0$. (B) Fraction of proteins bound to DNA as a function of $K_\theta$ under the same conditions as in panel (A). (C) Average $R_g$ of DNA as a function of $K_\theta$ for $\lambda_{PD} = 3.5$ with three different homotypic strengths $\lambda_{PP}$. (D) Fraction of proteins bound to DNA as a function of $K_\theta$ under the same conditions as in panel (C).

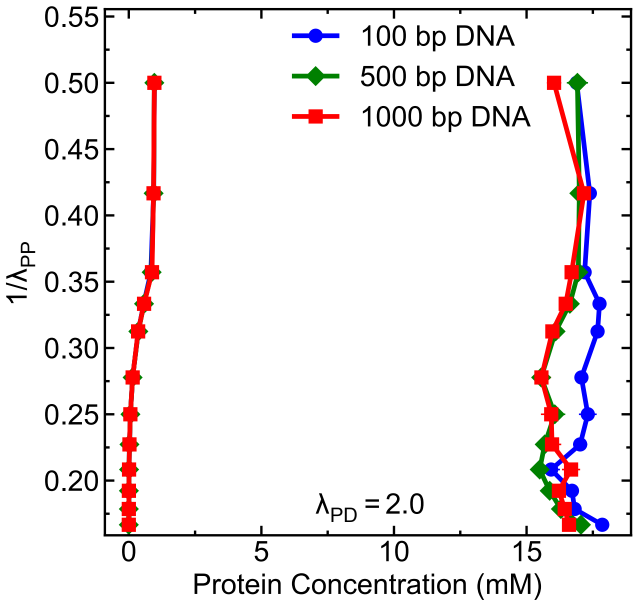


Figure S3: Phase diagram showing protein concentrations in the dilute and dense phases as a function of protein–protein interaction strength ($\lambda_{PP}$), for three different DNA lengths: 100 bp, 500 bp and 1000 bp at strong heterotypic interaction strength ($\lambda_{PD} = 2.0$).

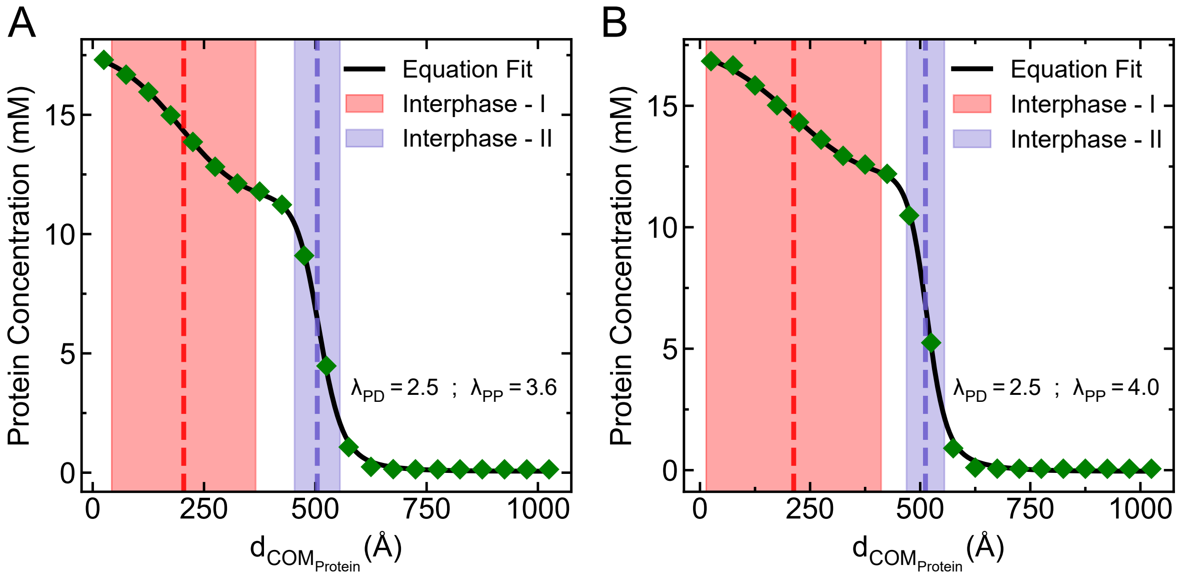


Figure S4: Radial density profile of proteins from the center of mass (COM) of proteins in the largest cluster at (A) $\lambda_{PD} = 2.5$, $\lambda_{PP} = 3.6$, and (B) $\lambda_{PD} = 2.5$, $\lambda_{PP} = 4.0$, showing two distinct protein populations. The profile is fitted with a double hyperbolic tangent function (solid black line, Eq. (S5)), capturing the presence of an inner DNA-associated protein-rich region (dense phase-I) and an outer protein-rich region not in contact with DNA (dense phase-II). Red and blue dotted lines represent the location of the two interphases between dense phase-I, dense phase-II and between dense phase-II and dilute phase, respectively. The DNA length considered in this analysis is 250 bp.

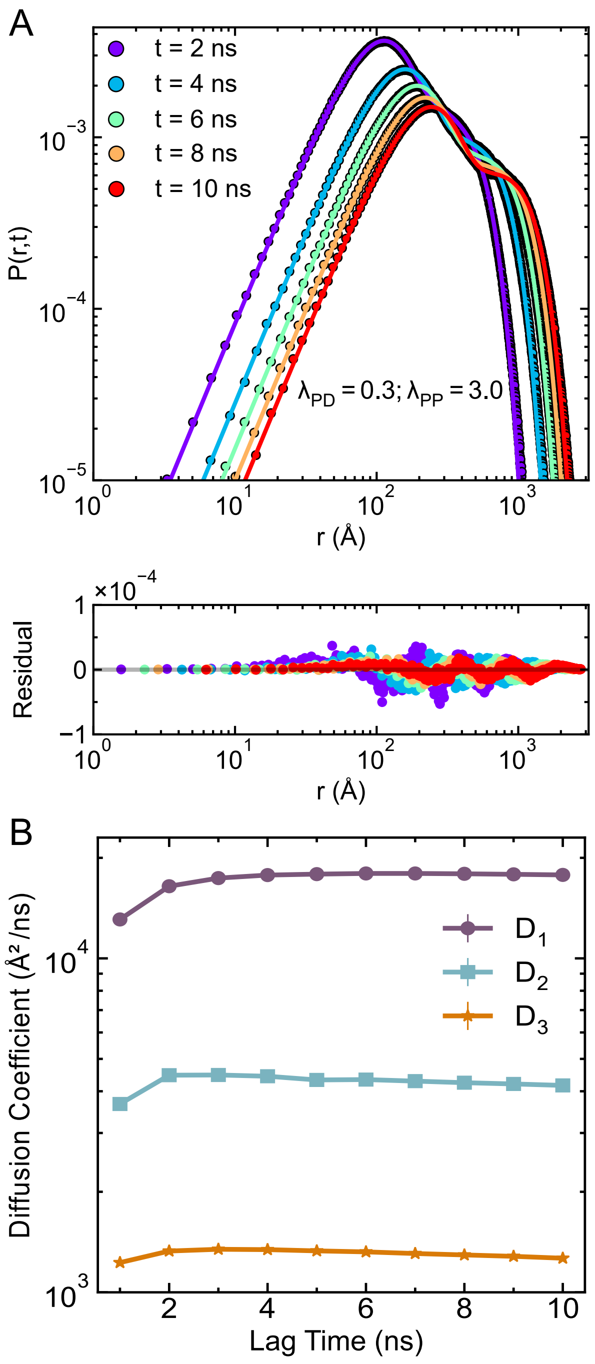


Figure S5: (A) Radial displacement probability distributions of proteins, $P(r, t)$, for different lag times $t$, under weak heterotypic ($\lambda_{PD} = 0.3$) and strong homotypic ($\lambda_{PP} = 3.0$) interaction strengths. The points denote histogram data from simulations, while solid lines show the best-fit using a sum of three Gaussian components (Eq. 6 in the main text). The difference between the Gaussian fitted lines and the simulation data points are represented as residuals from the data fitting for each lag time $t$ (lower panel), confirming the quality of fit. (B) Diffusion coefficients ($D_i$) corresponding to the three dynamic modes extracted from the Gaussian fits in (A), shown as a function of lag time. The plateau behavior indicates that each mode exhibits stable diffusivity across lag times.

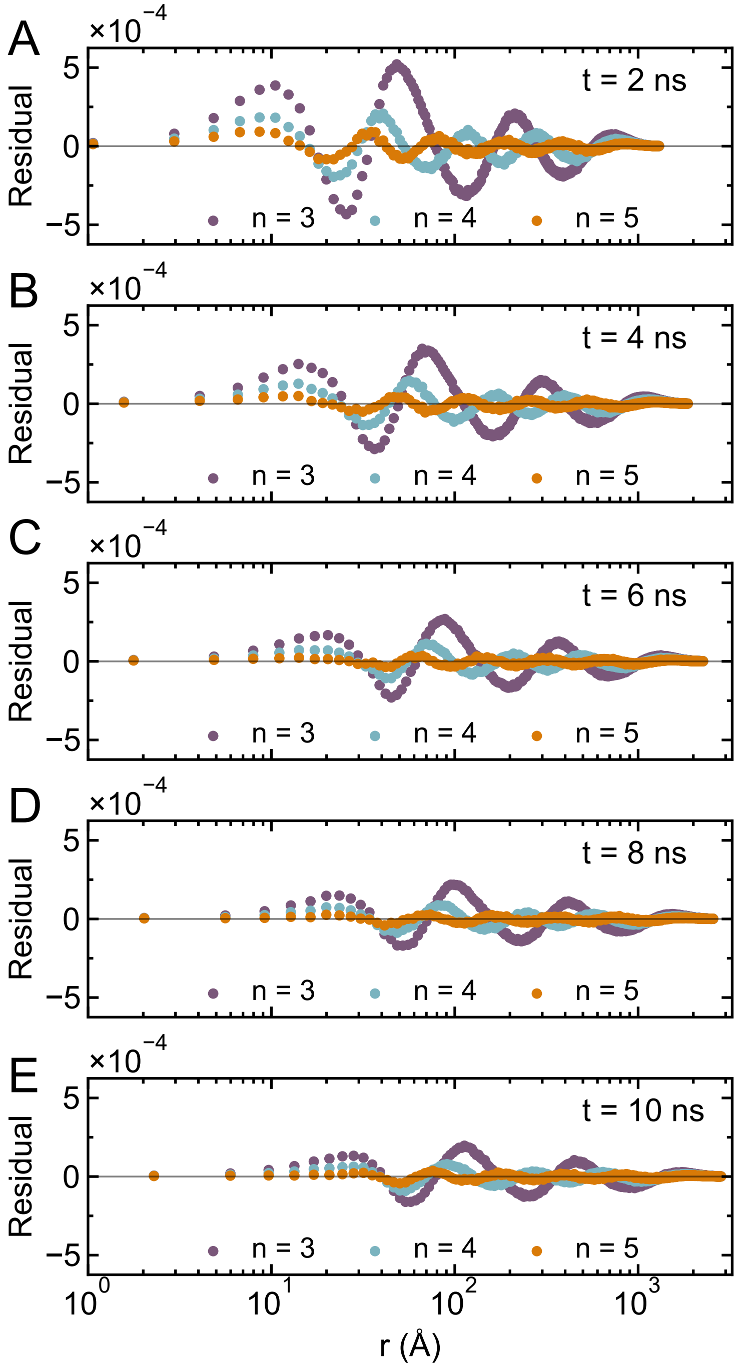


Figure S6: Residual analysis for Gaussian fitting of protein displacement distribution in multiphase condensates. To determine the optimal number of Gaussian components describing protein displacement distributions for the multiphasic condensate ($\lambda_{PD} = 2.0$ and $\lambda_{PP} = 3.0$ as considered in Fig. 7A in the main text), we calculated the residuals between simulation data and fitted curves using $n = 3$, $n = 4$, and $n = 5$ Gaussian components at a fixed lag time of (A) $t = 2$ ns, (B) $t = 4$ ns, (C) $t = 6$ ns, (D) $t = 8$ ns and (E) $t = 10$ ns, as a function of radial distance. The plot shows that the residual is minimized for $n = 5$ components at any lag time, confirming that a five-Gaussian model provides the best fit to the data across $r$.

### References


(1) Michaud-Agrawal, N.; Denning, E. J.; Woolf, T. B.; Beckstein, O. MDAnalysis: a toolkit for the analysis of molecular dynamics simulations. *Journal of computational chemistry* **2011**, *32*, 2319–2327.

(2) Devarajan, D. S.; Rekhi, S.; Nikoubashman, A.; Kim, Y. C.; Howard, M. P.; Mittal, J. Effect of charge distribution on the dynamics of polyampholytic disordered proteins. *Macromolecules* **2022**, *55*, 8987–8997.

(3) Ramasubramani, V.; Dice, B. D.; Harper, E. S.; Spellings, M. P.; Anderson, J. A.; Glotzer, S. C. freud: A software suite for high throughput analysis of particle simulation data. *Computer Physics Communications* **2020**, *254*, 107275.

(4) Kuo, I. W.; Mundy, C. J.; Eggimann, B. L.; McGrath, M. J.; Siepmann, J. I.; Chen, B.; Vieceli, J.; Tobias, D. J. Structure and dynamics of the aqueous liquid-vapor interface: a comprehensive particle-based simulation study. *The Journal of Physical Chemistry B* **2006**, *110*, 3738–3746.
